# dsRNA-based viromics: A novel tool unveiled hidden soil viral diversity and richness

**DOI:** 10.1101/2023.05.10.540251

**Authors:** A. Poursalavati, A. Larafa, M.L. Fall

## Abstract

Viruses play a crucial role in agroecosystem functioning. However, few studies have examined the diversity of the soil virome, especially when it comes to RNA viruses. Despite the great progress in viral metagenomics and metatranscriptomics (metaviromics) toward RNA viruses characterization, soil RNA viruses’ ecology is embryonic compared to DNA viruses. We currently lack a wet lab. method to accurately unhide the true soil viral diversity. To overcome this limitation, we developed dsRNA-based methods capitalizing on our expertise in soil RNA extraction and dsRNA extraction ported from studies of phyllosphere viral diversity. This proposed method detected both RNA and DNA viruses and is proven to capture a greater soil virus diversity than existing methods, virion-associated nucleic enrichment, and metaviromics. Indeed, using this method we detected 284 novel RNA-dependent RNA polymerases and expanded the diversity of *Birnaviridae* and *Retroviridae* viral families to agricultural soil, which, to our knowledge, have never been reported in such ecosystem. The dsRNA-based method is cost-effective in terms of affordability and requirements for data processing, facilitating large-scale and high-throughput soil sample processing to unlock the potential of the soil virome and its impact on biogeochemical processes (e.g. carbon and nutrient cycling). This method can also benefit future studies of viruses in complex environments, for example, to characterize RNA viruses in the human gut or aquatic environment where RNA viruses are less studied mainly because of technical limitations.

## Introduction

All cellular life forms are likely infected by viruses, which are the most ubiquitous and diverse biological entities on earth^1^. The soil is one of the highest viral-concentrated environments^2^. Through horizontal gene transfer, soil viruses influence microbial communities and act as a significant genetic resource that promotes biological diversity and evolution^3^.

The soil virome is important in ecology but remains inadequately characterized due to the difficulty in isolating viruses and their nucleic acids^2^. Soil micro-heterogeneity, matrix complexity, and organic inhibitors pose significant challenges to molecular biology techniques toward tapping soil viral diversity and abundance^4^. Metagenomics and microscopy-based studies have found DNA viruses to be the most abundant in soils^5–9^. However, current soil virus identification methods encompass a systematic bias toward prokaryotic viruses, particularly encapsulated viruses, to the detriment of eukaryotic viruses that lack capsid formation^10^. As a result, soil viral ecology research has mainly focused on DNA viruses of bacteria and archaea.

Given the possible host diversity in soil, it is expected that a large number of RNA viruses may exist. Most of the research on soil RNA viruses has focused on single hosts for important crops or crop diseases^15^ ^16^. Generating and analyzing environmental RNA viral sequencing data is challenging due to technical limitations in extracting enough viral RNA from complex biotope samples and computationally finding viral genomic signature in large metatranscriptomic datasets^13^. To overcome these limitations associated capturing soil RNA viruses, a recent study has modified the VANA method to capture RNA viruses by eliminating the RNA digestion step and sequencing all extracted RNA including those from dominant organisms^10^. The authors were able to shown a divers community of soil RNA viruses, including viruses from bacteria, plants, fungi, vertebrates, and invertebrates, which give a novel insights on soil viral diversity.

Historically, two soil viral genetic material extraction methods has been widely used for detection of soil viruses through sequencing, Virion-Associated Nucleic Acids (VANA) and total RNA or DNA extractions. The VANA method is highly specific to viral species detection, but encompass a bias toward DNA viruses^11^. The second method is referred to metaviromics (metagenomics and metatranscriptomics applied to viruses)^12^. The main challenges with use of soil metaviromics is that it relies on sequencing genetic materials dominated by organisms with larger genomes (e.g. bacteria, fungi etc.), making difficult to accurately identify viruses^10, 13^. Furthermore, metaviromics is expensive, error-prone and time-consuming when trying to identify viruses^14^. In addition, these methods for detecting soil viruses (VANA and metaviromics) can detect either DNA or RNA viruses at the same time and may introduce biases when characterizing the soil virus diversity.

Over the last five years, numerous metaviromics studies has help to expand soil RNA viral diversity and ecology^11, 17, 18^. The majority of soil RNA viruses have been classified using the RNA-dependent RNA polymerase (RdRp) as a phylogenetic marker^19^. This technique has been used to significantly increase the number of known RNA viral sequences in divers ecosystems^20–24^. The RdRp plays a critical role in RNA virus genome replication and transcription, and is therefore widely used as a reference gene to estimate RNA virus phylogenies, forming the basis for the newly established classification scheme for RNA viruses^25^. The RdRp domain sequences contain eight conserved amino acid motifs, with the motif A, B and C being highly characterized^26^. The RdRp approach is a unique and novel way of studying RNA viruses, made possible by the development of advanced tools and databases that allow for efficient analysis of large datasets^27^. In recent years, several new tools and databases have emerged and greatly facilitated the use of the RdRp approach in virology research, including the NeoRdRp database^28^ and the Muscle5 software package^29^. These tools and databases enable the identification of conserved RdRp domains and motifs and the construction of robust phylogenetic trees^30^. Despite these bioinformatics-based advanced tools, the effects of soil RNA viruses on soil ecological processes are still mostly unknown because of forementioned metaviromics limitations and the lack of viromic-based wet lab method to upscale virus ecology from soil ecoystems^31^.

Most, if not all, the viruses generate double-stranded RNA (dsRNA) as an intermediate during the viral replication process or as an erroneous product resulting from the bidirectional transcriptional readthrough^32–35^. Consequently all viral hosts have conserved mechanisms by which they sense viral infections through detection of dsRNA^34^. Therefore, the dsRNA molecule can serve as a stable and resistant matrix for virus characterization, due to its thermodynamic stability and resistance to RNase destruction^36^, especially in situations where virus purification is challenging due to low titer, particle instability, or the presence of inhibitors. The extraction of dsRNA is a common approach widely used for the characterization of plant and fungi virome^37–40^. Gaafar and Ziebell in 2020^37^ conducted a comparative study between viral particle extraction and dsRNA extraction in plants and concluded that the dsRNA-based approach should be prioritized due to its consistent ability to provide a more thorough understanding of the virome. However, to the best of our knowledge, there have been no published attempts to employ the dsRNA-based approach for soil virome characterization thus far.

In this study, a two-fold objective was pursued. Firstly, to enhance the detection and classification of viruses in organic and mineral soils through dsRNA-based methods, highlighting the significance of soil types and extraction method in analyzing RNA viral communities and their ecological roles. A cost-effective dsRNA-based method was developed and shown to expand the diversity of viral families in agricultural soil. Secondly, the investigation aimed to explore the soil virome’s diversity through dsRNA-based approaches by identifying RdRp regions in assembled sequences. Different extraction methods showed distinct viral community compositions, with mineral soils having more eukaryotic viruses and organic soils having a more diverse community, including dsDNA viruses.

## Results

For the sake of convenience and clarity, all the results (figures and tables) related to organic soil were put in the supplementary material section and can be easily consulted.

### Bioinformatics and data processing

The sequencing results obtained from the three extraction methods (CCC-dsRNA, RPT-dsRNA, and VANA), were presented in table 1 and included the average number of reads, average number of bases, and average read length (after trimming) (Table 1 and Supplementary File 1). The results indicated that dsRNA-based methods produced both the highest average number of reads and the highest average number of bases. On the other hand, VANA generated the longest average read length.

**Table 1:**
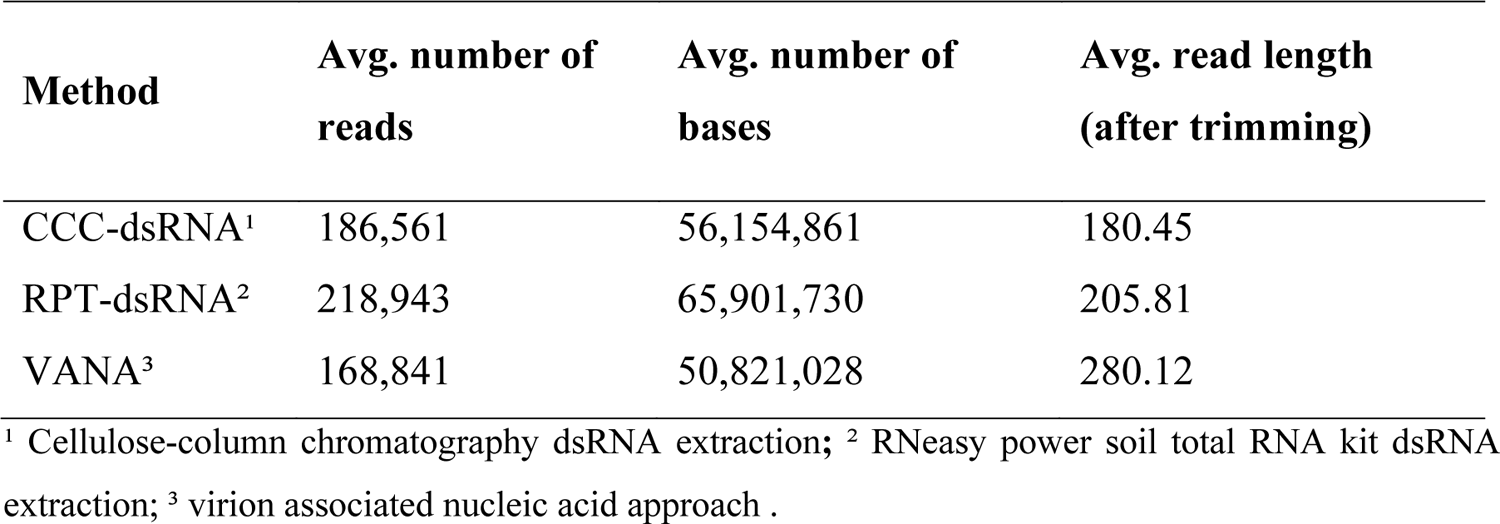
Reads statistics for sequencing run of the three extraction methods

| Method | Avg. number of reads | Avg. number of bases | Avg. read length (after trimming) |
| --- | --- | --- | --- |
| CCC-dsRNA<sup>1</sup> | 186,561 | 56,154,861 | 180.45 |
| RPT-dsRNA<sup>2</sup> | 218,943 | 65,901,730 | 205.81 |
| VANA<sup>3</sup> | 168,841 | 50,821,028 | 280.12 |

<sup>1</sup> Cellulose-column chromatography dsRNA extraction; <sup>2</sup> RNeasy power soil total RNA kit dsRNA extraction; <sup>3</sup> virion associated nucleic acid approach .

To analyse our sequencing data, a new bioinformatics workflow was developed in Python language and compiled in a pipeline named SOVAP (Soil Virome Analysis Pipeline)^41^, which we described in the material and methods section. The pipeline comprises several modules that were designed to process the raw sequencing data and to identify, classify, and quantify the virome present in the samples (Fig. 1).

**Figure 1:**
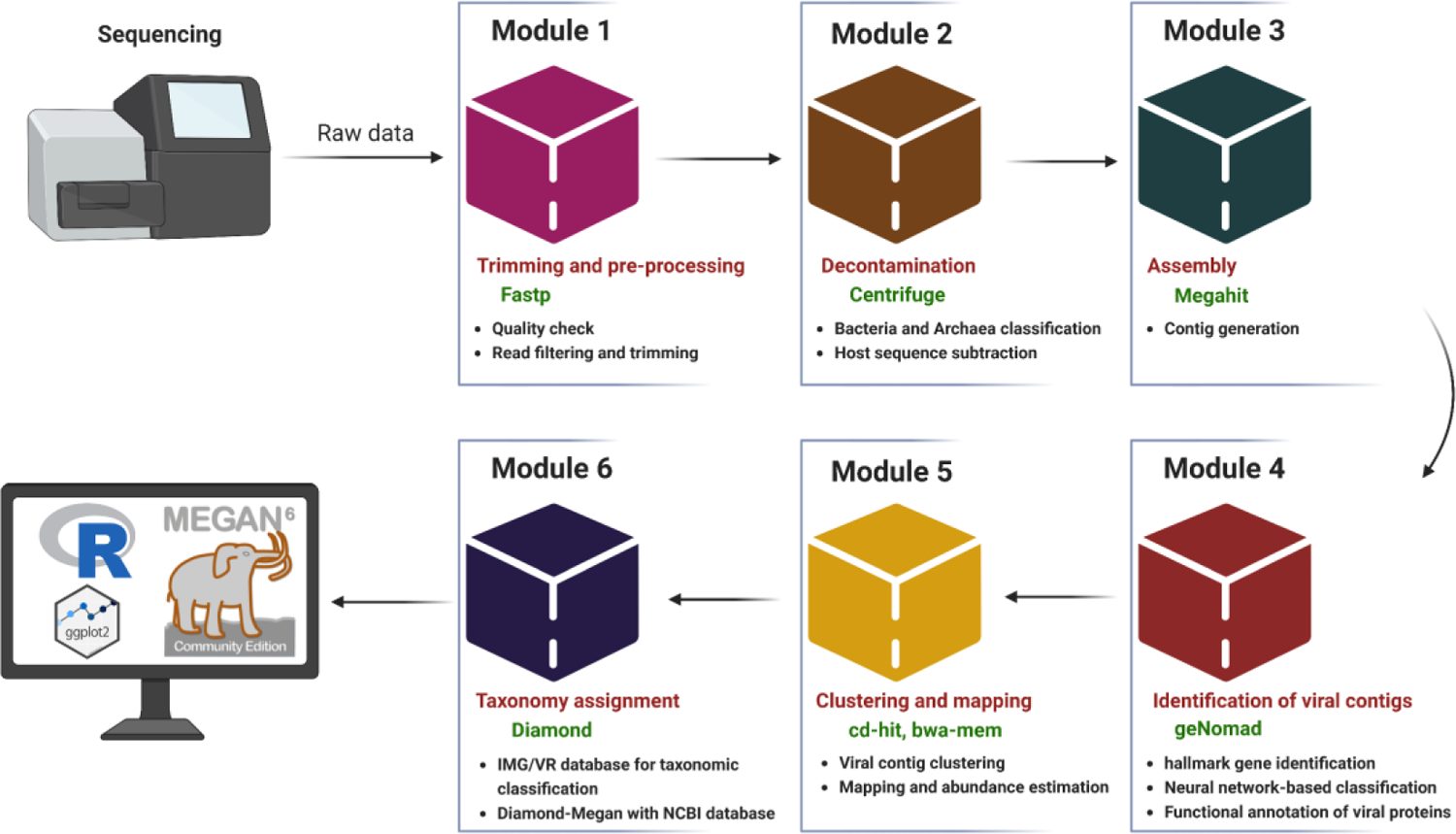
Overview of the SOVAP (soil virome analysis pipeline) workflow for viromics data processing and analysis. Module 1 includes quality checking and trimming of raw sequencing reads using Fastp. In module 2, contaminants are removed using Centrifuge. Clean reads are then assembled into contigs in module 3 using Megahit. In module 4, viral contigs are identified with the help of geNomad. In module 5 the abundance of viral contigs is estimated and redundant contigs are clustered. Module 6 involves the annotation and taxonomy assignment of viral contigs using Diamond and IMG/VR database.

In the second module of the SOVAP pipeline, which is dedicated to the decontamination step, we used Centrifuge software to filter out reads from non-viral sources. To achieve this, we created a custom database for Centrifuge which derived from SARs, Nematodes, Bacteria, Archaea, Fungi, Protozoa, and Human reference and representative genomes obtained from NCBI RefSeq (access date 16.03.2023) using Genome_updater v0.5.1 (https://github.com/pirovc/genome_updater) (which includes 4,668 genomes)^42^. This custom database enables Centrifuge to accurately identify and remove non-viral reads from the virome datasets, resulting in more accurate and reliable viral detection and analysis. As part of the sixth module of the pipeline, a Diamond index of the Genbank viral database containing 58,201 genomes (access date 18.03.2023) is provided. This database is used in the last module when the IMG/VR database is not available or the pipeline is running in Diamond-Megan mode.

### Viral taxonomy

The analysis of viral communities in mineral and organic soils using three different extraction methods yielded distinct taxonomic insights at various taxonomic ranks (Supplementary Files 2 and 3). The results showed that the VANA method displayed a viral communities dominated by DNA viruses, particularly *Caudoviricetes*, a type of tailed bacteriophages, which represented 92-94% of the TPM (Fig. 2 and Supplementary Fig. 1).

**Figure 2:**
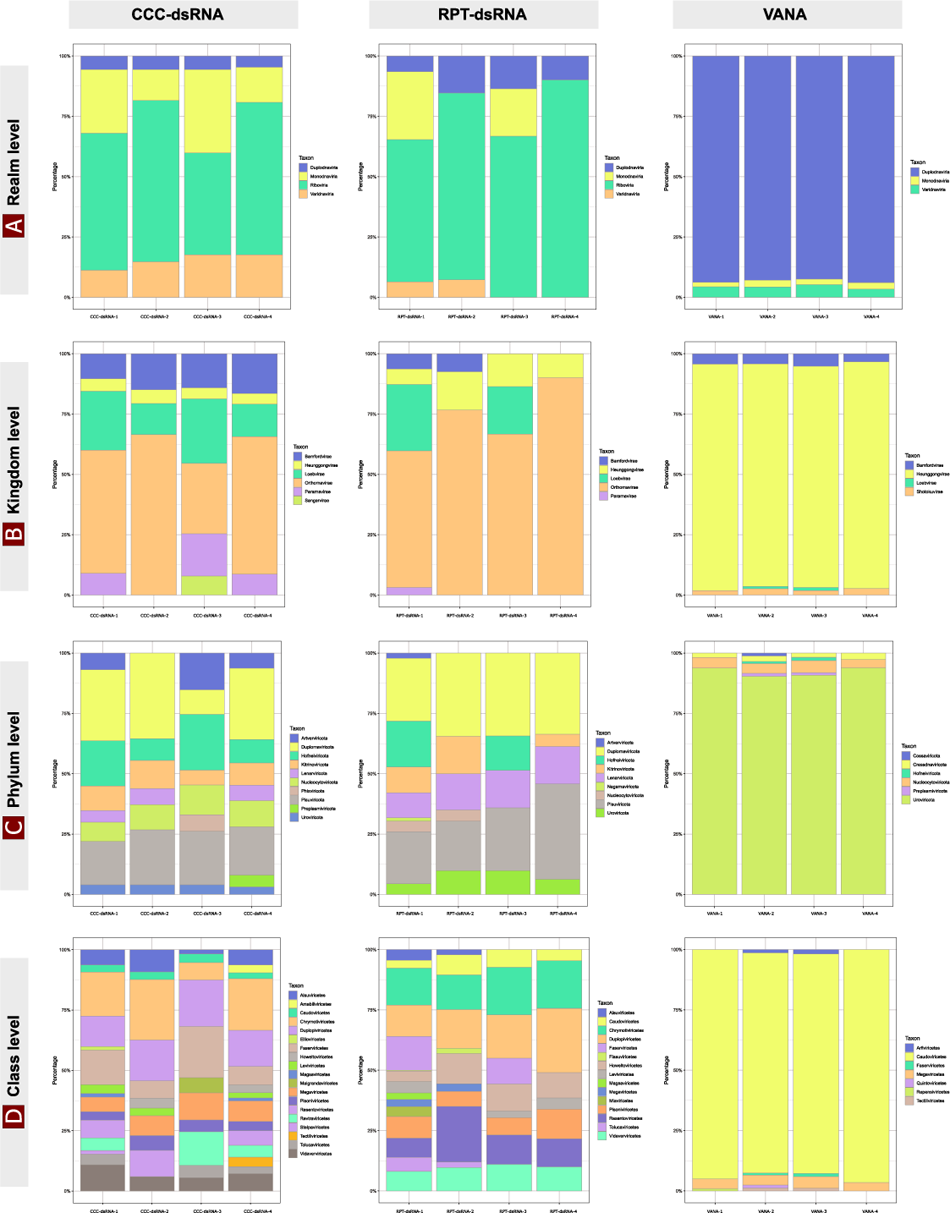
Comparison of taxonomic composition of viral communities in four mineral soil sample replicates extracted using three different methods: RNeasy Power Soil Total RNA kit (RPT-dsRNA), Cellulose-Column Chromatography (CCC-dsRNA), and the Virion Associated Nucleic Acid approach (VANA), at the taxonomic ranks of (a) realm, (b) kingdom, (c) phylum and (d) class. The stacked bar chart displays the percentage of assigned viral sequences for each taxonomic level.

In mineral soil samples, the dsRNA-CCC and dsRNA-RPT extractions showed similar viral community compositions, with the majority of the viral contigs belonging to the *Pisuviricota* and *Duplornaviricota* phyla, which include both +ssRNA and dsRNA viruses. The *Uroviricota* phylum was also present in both extractions but was less abundant in the dsRNA methods than in the VANA extraction. The *Negarnaviricota* phylum, which comprises -ssRNA eukaryotic viruses, as well as *Kitrinoviricota* and *Lenarviricota*, which include +ssRNA eukaryotic and prokaryotic viruses, respectively, were also detected (Fig. 2, and Supplementary Fig. 3). Only the CCC-dsRNA extraction method was able to detect the presence of *Phixiviricota* phylum (non-enveloped ssDNA viruses) in this particular soil type (Fig. 2, and Supplementary Fig. 3). In contrast, the VANA extraction method yielded a different viral community composition in mineral soil samples, with the majority of the viral contigs belonging to the *Uroviricota* phylum, which includes tailed bacteriophages that infect bacteria. Additionally, *Nucleocytoviricota*, which includes dsDNA eukaryotic viruses, and *Cressdnaviricota*, which includes ssDNA viruses, were also present (Fig. 2, and Supplementary Fig. 3).

At the family level, the dsRNA methods were dominated by *Totiviridae* and *Partitiviridae*, both including dsRNA viruses. *Mitoviridae* and *Tombusviridae*, which include +ssRNA eukaryotic RNA viruses and +ssRNA plant viruses, respectively, were present in both dsRNA extraction methods but were more abundant in the RPT-dsRNA extraction method. Both dsRNA methods also detected *Retroviridae*, a family of single-stranded RNA viruses that infect vertebrates. However*, Birnaviridae*, which includes dsRNA viruses, and *Virgaviridae*, which includes +ssRNA viruses, were only present in the RPT-dsRNA method (Fig. 3, and Supplementary Fig. 3).

**Figure 3:**
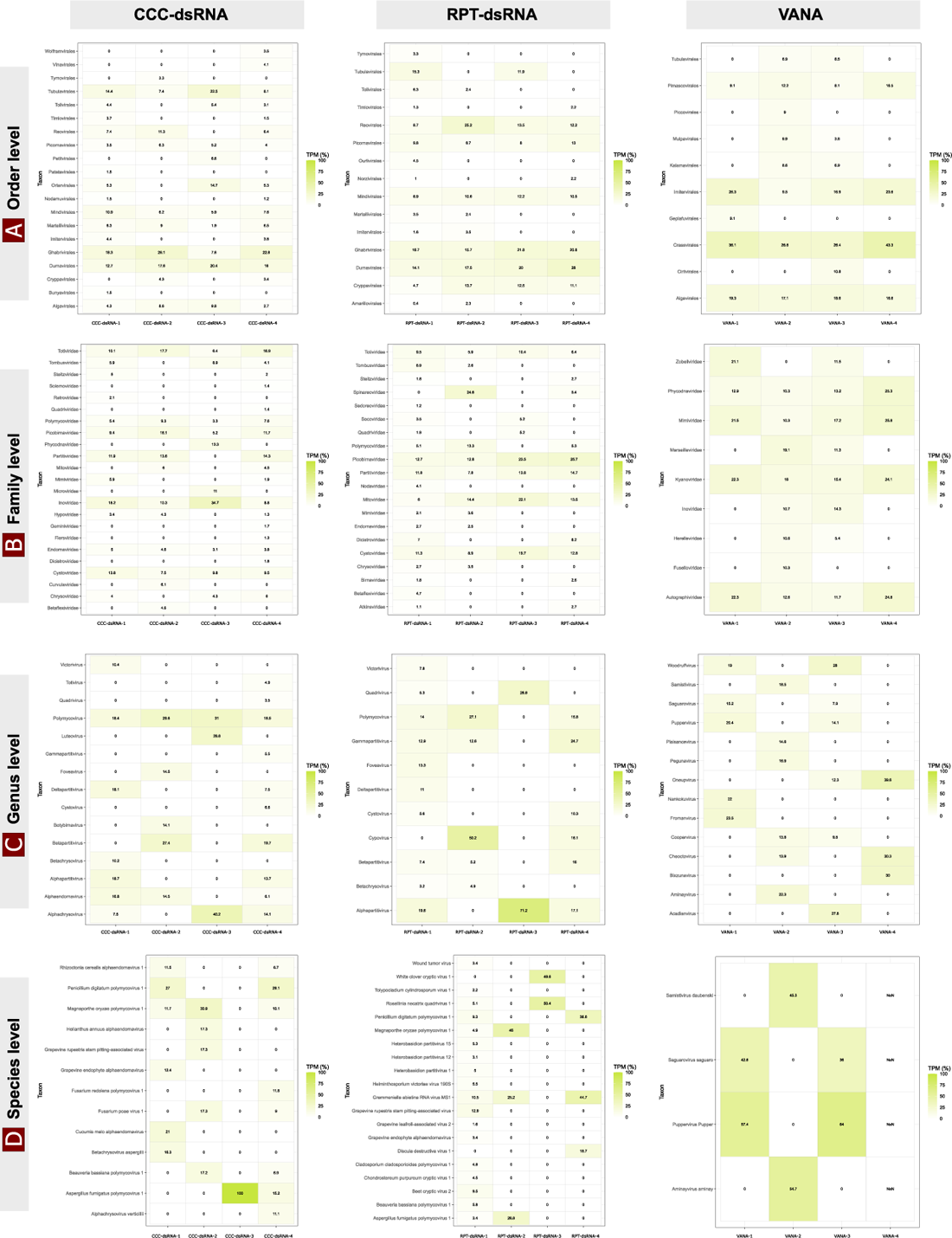
Comparison of viral taxa abundance across four mineral soil sample replicates using three different extraction methods: RNeasy Power Soil Total RNA kit (RPT-dsRNA) and Cellulose-Column Chromatography (CCC-dsRNA), and the Virion Associated Nucleic Acid approach (VANA) at the (a) Order, (b) Family, (c) Genus and (d) Species levels. A heatmap shows the relative abundances determined by calculating the Transcripts per million (TPM) value of the reads counts of each viral taxa and their relative abundances in each sample replicate. The percentages of each viral taxa are indicated in the corresponding rectangles.

In organic soil samples, viruses of *Duplornaviricota* phylum were the most abundant in the RPT-dsRNA extraction method, while with the CCC-dsRNA method viruses of *Artverviricota* phylum were the most abundant, including all ssRNA-RT and dsDNA-RT viruses (Supplementary Fig. 1). The VANA approach revealed *Uroviricota* as the most abundant phylum and the presence of *Preplasmiviricota* phylum. At the family level when extracted using the VANA approach, the families *Tectiviridae*, *Straboviridae*, and *Demerecviridae*, which include dsDNA viruses, were more abundant. The *Inoviridae* family of ssDNA viruses and even *Mimiviridae* giant virus were observed not only in VANA but also in the CCC-dsRNA extraction (Supplementary Figs. 2 and 4). The RPT-dsRNA extraction method gave a more diverse viral community than the CCC-dsRNA method. Indeed, the RPT-dsRNA extraction method showed a different viral community composition, with the majority of the viral contigs belonging to the family *Cistroviridae*, dsRNA viruses that infect bacteria. Also unique families were observed using the RPT-dsRNA extraction method, including *Retroviridae* (+ssRNA viruses) and *Partiviridae* (dsRNA viruses) (Supplementary Figs. 2 and 4). The CCC-dsRNA extraction method revealed the presence of *Tombusviridae* (+ssRNA of plant viruses), *Phycodnaviridae* (dsDNA eukaryotic viruses). *Birnaviridae* and *Totiviridae* as well as phylum of *Phixiviricota* (non-enveloped ssDNA viruses) were detected with both RPT-dsRNA and CCC-dsRNA methods (Supplementary Figs. 2 and 4).

**Figure 4:**
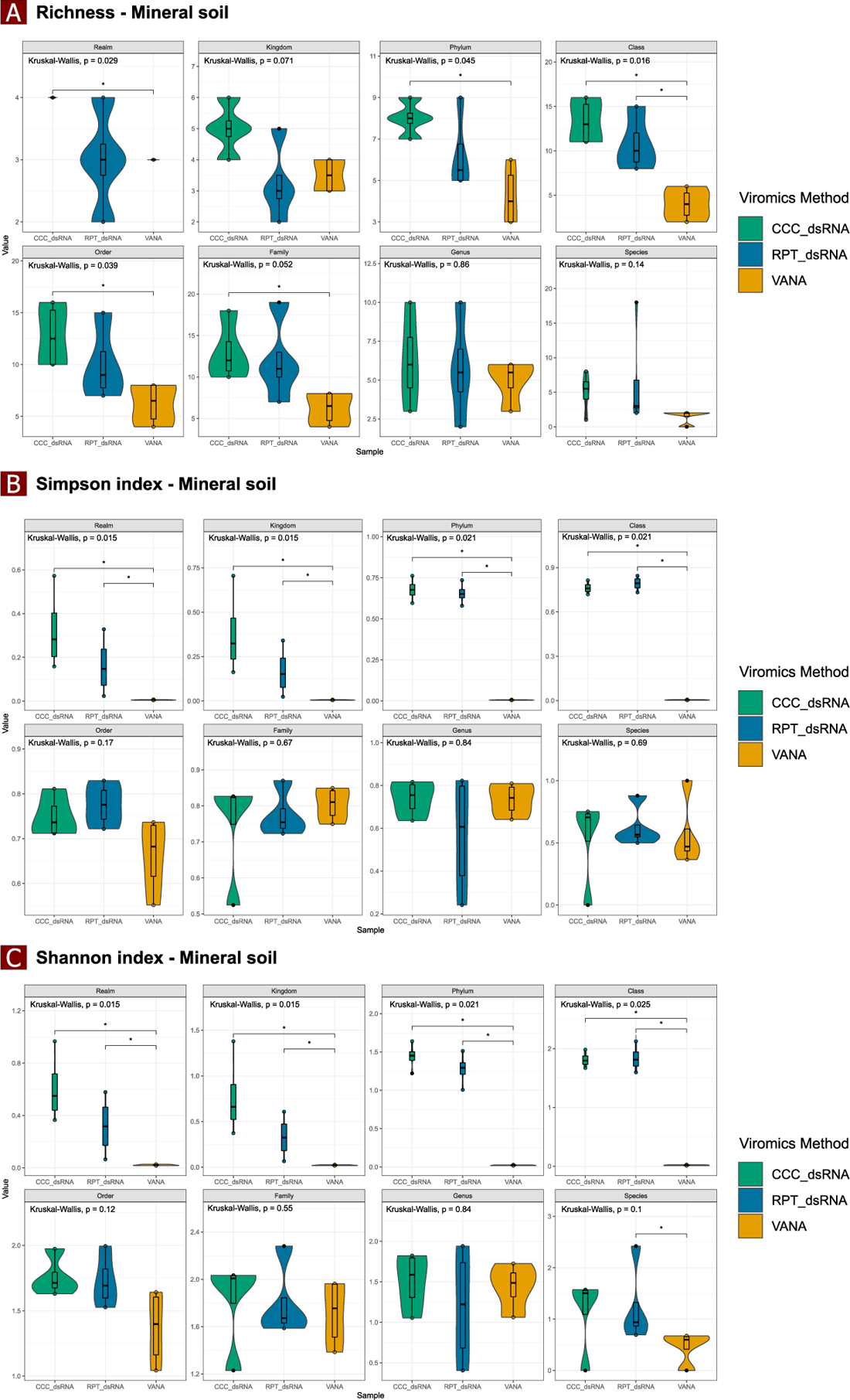
Alpha-diversity analysis of viral community composition in mineral soil using three extraction methods: RNeasy Power Soil Total RNA kit (RPT-dsRNA) and cellulose-column chromatography (CCC-dsRNA), and the Virion Associated Nucleic Acid approach (VANA). Violin plots show the Richness (A), Shannon (B), and Simpson (C) indices of all biological replicates of each method, calculated using Vegan v2.6-4 and Tidyverse packages. The analysis was performed at the realm, kingdom, phylum, class, order, family, genus, and species levels.

## Viral diversity

### Alpha diversity

The differences in the alpha diversity of viral DNA and RNA viruses using the three different virome enrichment methods: CCC-dsRNA, RPT-dsRNA, and VANA were also investigated. The richness, Shannon, and Simpson indices at different levels of taxonomic rank (realm, kingdom, phylum, class, order, family, genus, and species) showed significant (P < 0.05) differences in alpha diversity between the three extraction methods in both mineral and organic soil samples. Specifically, in mineral soil, the Simpson diversity index was significantly different at the realm (p = 0.015), kingdom (p = 0.015), phylum (p = 0.021), and class (p = 0.021) levels, while no significant differences (P > 0.05) were observed between the methods at the order, family, genus, and species levels (Fig. 4b and Supplementary Fig. 6b). The richness index was significantly different between the three extraction methods at the realm (p = 0.029), phylum (p = 0.045), class (p = 0.016), and order (p = 0.039) levels, with the CCC-dsRNA extraction method resulting in the highest index (Fig. 4a and Supplementary Fig. 6a). In organic soil, the Simpson index was significantly different between the three extraction methods, with the highest index recorded at the realm (p = 0.0097), kingdom (p = 0.0073), phylum (p = 0.0073), and class (p = 0.024) levels with the dsRNA extraction methods. At the family level (p = 0.025), the VANA extraction approach recorded the highest index (Supplementary Figs. 5b and 7b). The Shannon index was significantly higher in mineral soil in both dsRNA-based methods at the realm (p = 0.015), phylum (p = 0.015), kingdom (p = 0.021), and class (p = 0.025) levels (Fig. 4c and Supplementary Fig. 6c), while it was higher in organic soil in the VANA method at the family (p = 0.023), genus (p = 0.009), and species (p = 0.0092) levels, and in the dsRNA extraction methods at the realm (p = 0.097), kingdom (p = 0.073), phylum (p = 0.018), and class (p = 0.018) levels (Supplementary Figs. 5c and 7c).

### Beta diversity

Using the Bray-Curtis dissimilarity metric, Principal Coordinates Analysis (PCoA) was performed to compare viral communities in the two soil types extracted using the three different methods: CCC-dsRNA, RPT-dsRNA, and VANA. The beta diversity analysis showed distinct results for the mineral and organic soil samples. In mineral soil, the viral communities in the CCC-dsRNA and RPT-dsRNA methods clustered more closely compared to the viral community resulting from VANA extraction, indicating a more similar community composition. The viral community resulting from VANA method was clearly separated from the CCC-dsRNA and RPT-dsRNA samples at all taxonomical levels, suggesting a unique viral community (Fig. 5). In organic soil, the viral community resulting from the CCC-dsRNA and RPT-dsRNA methods not only clustered more closely, but interestingly there were shared viral community between the these two methods. In contrast, the viral community resulting from the VANA method was clearly separated from the viral community resulting from the dsRNA-based methods at the realm, kingdom, and phylum levels. The CCC-dsRNA, RPT-dsRNA, and VANA viral communities were also clustered separately at the class, order, and family levels, indicating distinct microbial communities at these taxonomic levels (Supplementary Fig. 8). The PERMANOVA results showed that the differences observed in the PCoA were significant (p < 0.05) at the different taxonomic levels for both soil types (Fig. 5 and Supplementary Fig. 8).

**Figure 5:**
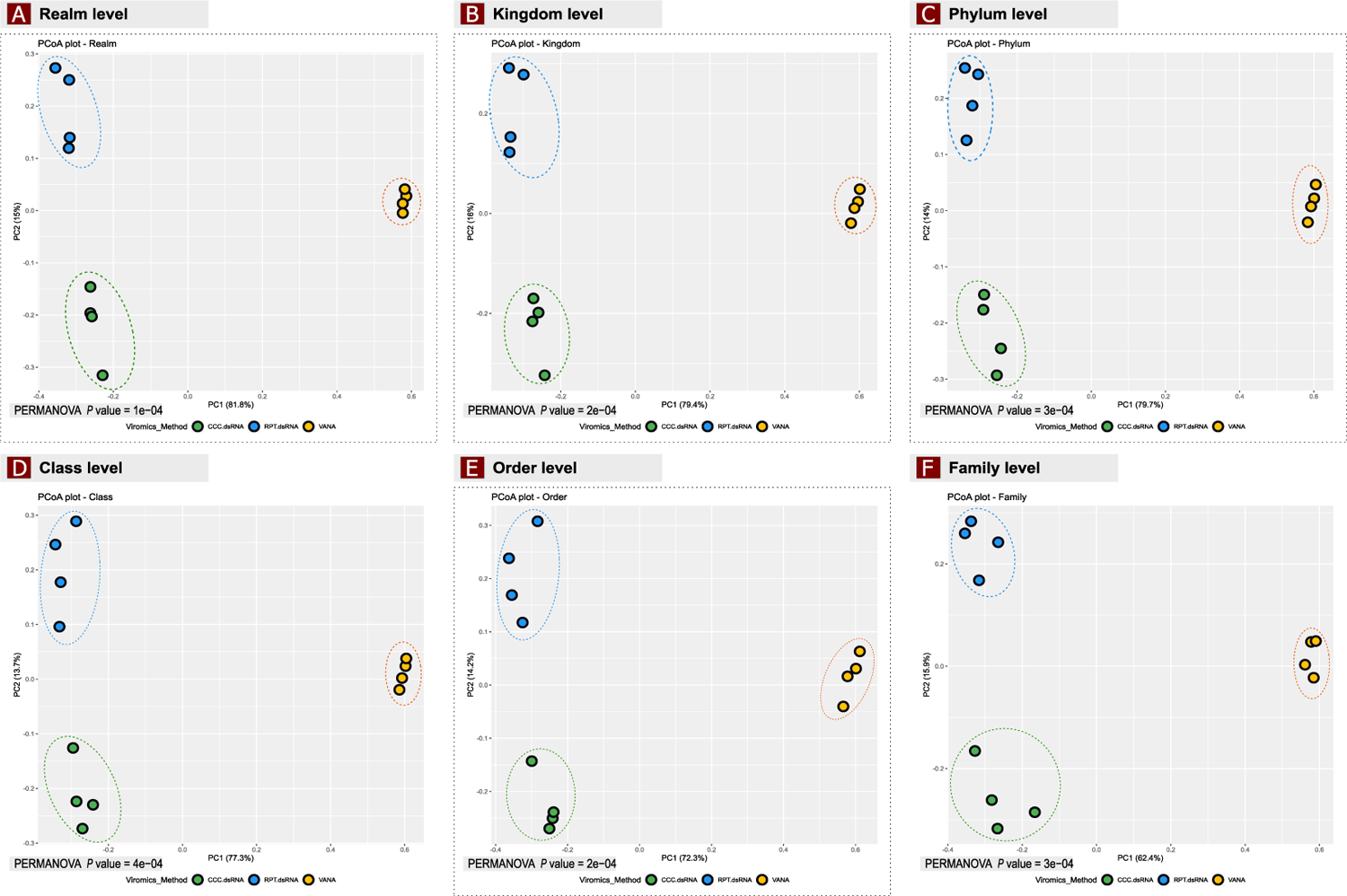
Principal Coordinates Analysis (PCoA) plot of viral community composition in mineral soil, comparing the effectiveness of three extraction methods: RNeasy Power Soil Total RNA kit (RPT-dsRNA), Cellulose-Column Chromatography (CCC-dsRNA), and Virion Associated Nucleic Acid approach (VANA), based on pairwise Bray-Curtis dissimilarities at each taxonomic rank. Each point on the plot represents a soil sample colored by the extraction method used. The p-value indicates the statistical significance of differences in viral community composition, as determined by a Permutational multivariate analysis of variance (PERMANOVA) test.

### Viral phylogenetic analysis

In both soil types, the dsRNA-based methods yielded an average of 5000-26000 viral contigs per sample through dsRNA extractions (Supplementary File 1). Therefore, we focused on the identification of novel contigs in *Riboviria* taxa candidates using RNA-dependent RNA polymerase (RdRp) as a phylogenetic marker and key RdRp motifs A, B, and C multiple sequence alignments analysis (Figs. 6 and 7). In total, we identified 3814 RdRps using functional RdRp protein motif annotations obtained from the RdRp-Scan workflow and database^27^. Of these, 2944 were unique. Through multi-sequence alignment analysis of the conserved A, B, and C protein motifs of the RdRp domains (Fig. 6), we identified 284 potentially novel sequences that were classified into different phyla and family (Supplementary File 4). The *Duplornaviricota* (dsRNA viruses) phylum had the highest number of novel RdRps, with a total of 198 (Supplementary Fig. 9). We also identified 48 novel RdRps in the *Pisuviricota* phylum, which includes positive-sense single-stranded RNA and some double-stranded RNA viruses (Supplementary Fig. 10). In addition, we found 18 novel RdRps in the *Kitrinoviricota* phylum, which are positive-sense single-stranded RNA viruses that infect eukaryotes (Supplementary Fig. 11). We also identified 18 novel RdRps in the *Lenarviricota* phylum, which are positive-sense single-stranded RNA viruses that infect prokaryotes (Supplementary Fig. 12). Finally, we found two novel RdRps in the *Birnaviridae* family, which are dsRNA viruses (Supplementary Fig. 13).

**Figure 6:**
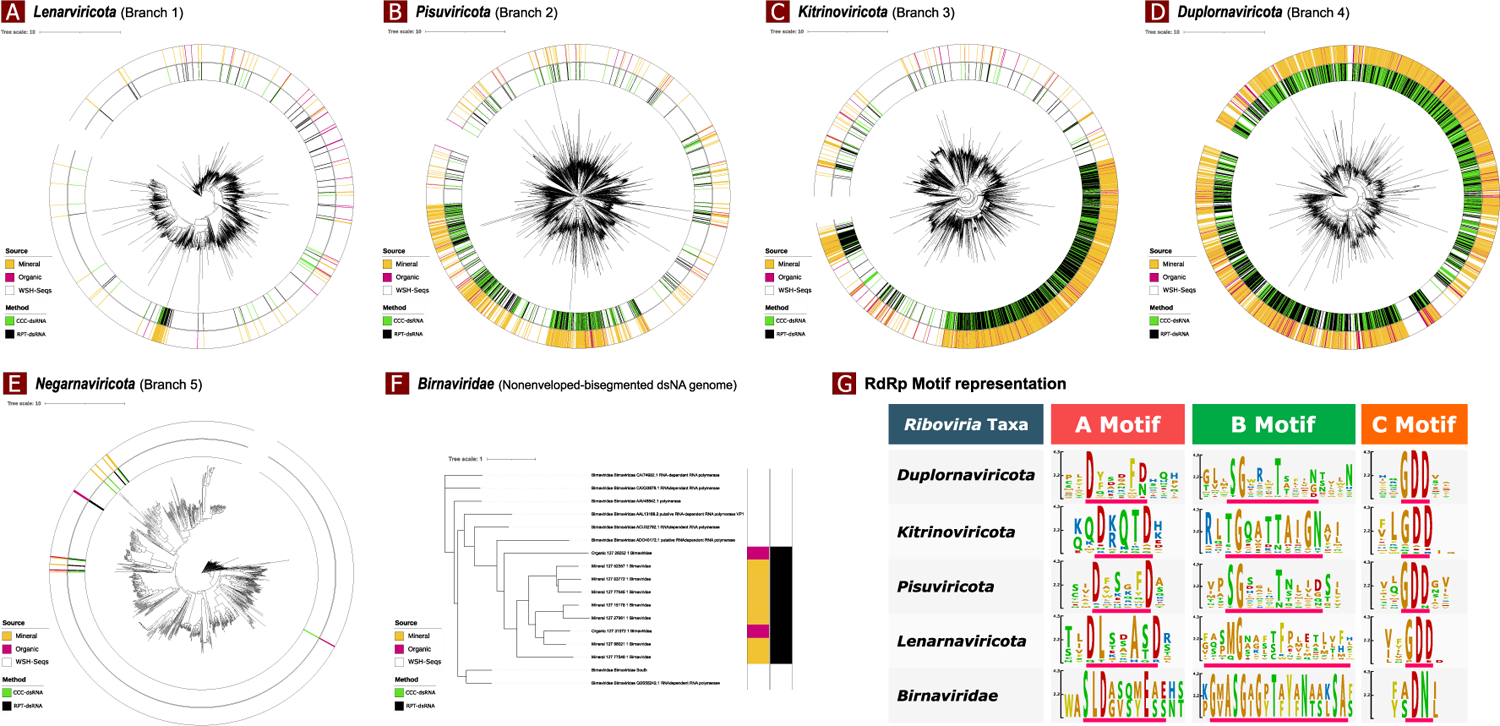
Phylogenetic trees of RdRp sequences. The RdRp sequences extracted from mineral (inner ring, yellow) and organic (inner ring, pink) soil types using two dsRNA extraction methods, Cellulose-Column Chromatography (CCC-dsRNA) (outer ring, green) and RNeasy Power Soil Total RNA kit (RPT-dsRNA) (outer ring, black), were aligned with those reported in previous studies (WSH; Wolf, Starr, and Hillary, inner ring, white) and published by Hillary. The trees are divided into the RNA viral phyla, including *Lenarviricota* (a), *Pisuviricota* (b), *Kitrinoviricota* (c), *Duplornaviricota* (d), *Negarnavirota* (e), and *Birnaviridae* Family (f). The conserved motifs logos representing each *Riboviria* taxa that were extracted from the multiple sequence alignment results are shown in (g).

**Figure 7:**
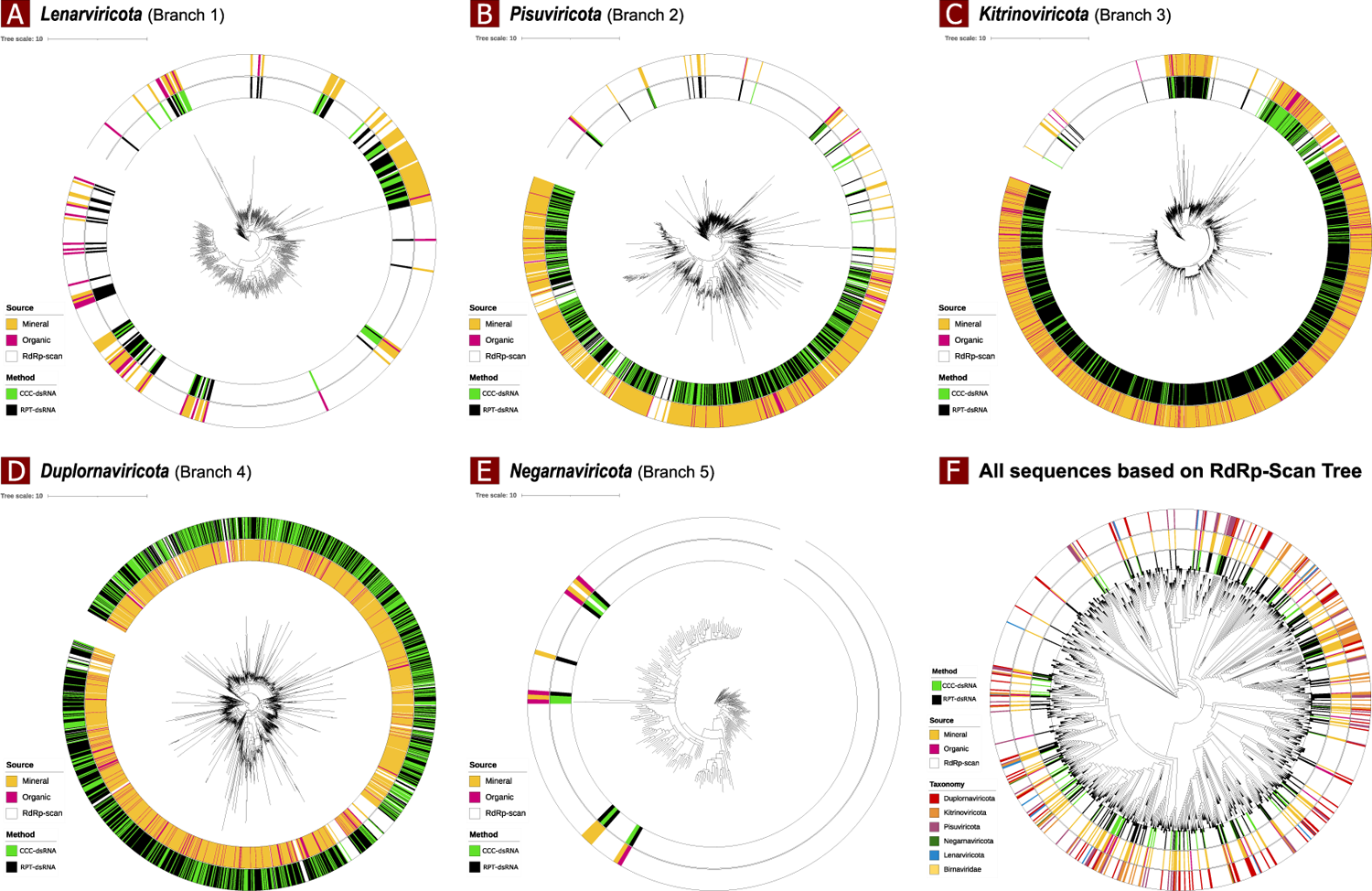
Phylogenetic trees of RdRp sequences. The RdRp sequences extracted from mineral (inner ring, yellow) and organic (inner ring, pink) soil types using two dsRNA extraction methods, Cellulose-Column Chromatography (CCC-dsRNA) (outer ring, green) and RNeasy Power Soil Total RNA kit (RPT-dsRNA) (outer ring, black), were aligned to the pre-built RdRp-Scan backbone trees published by Charon (inner ring white). The global RNA viral phylogeny is classified into the proposed RNA viral phyla, including (a) *Lenarviricota*, (b) *Pisuviricota,* (c) *Kitrinoviricota*, (d) *Duplornaviricota*, (e) *Negarnavirota*. (f) The diversity of RNA viruses across all identified taxa is visualized in a single phylogenetic tree using the RdRp-Scan whole tree backbone, providing a comprehensive view.

## Discussion

Viruses are incredibly abundant on Earth, with estimates suggesting that there could be 10^31^ viruses^17, 18, 44–46^. However, only 10^4^ viral species are currently listed in the International Committee on Taxonomy of Viruses (ICTV) official masters species list^47^. In addition, when it comes to soil environments, our understanding of viral diversity and interactions is limited to selected groups of viral families largely dominated by DNA viruses^44^. The difficulty and the cost in extracting soil RNA viruses has resulted in a slower discovery rate compared to dsDNA viruses^48^ ^13, 12^. In 2022, a novel metaviromics method (metagenomics and metatranscriptomics applied to viruses^12^) was able to detect soil RNA viruses, but had missed important RNA virus taxa and still an expensive method for massive soil samples processing^10^. The affordable and efficient viromics method proposed in the study is a known need for the scientific community to harness the soil viral diversity and fill the knowledge gaps on soil virus ecology, RNA viruses in particular.

The dsRNA-based methods captured a greater soil viral diversity than the standard method (VANA). When using the dsRNA-based methods compared to standard method, the alpha diversity was significantly higher at the four top ranks (real, kingdom, phylum and class) of virus taxonomy. The dsRNA-based methods in mineral soil yielded a significantly higher viral diversity and richness at the four and five top rank taxa, respectively. Indeed, with the VANA method the alpha diversity at all virus taxonomic levels was significantly lower (or similar) than dsRNA-based methods, except for organic soil where the alpha diversity obtained from the VANA method was significantly higher at the family and genus levels. However, the taxonomy analysis confirmed the known bias toward DNA viruses of the VANA method with 92-94% of the TPM belonging to the class of *Caudoviricetes*, dsDNA viruses known as tailed bacteriophages. Regardless of the soil types, VANA method wasn’t able to detect RNA viruses of *Riboviria* realm and missed 50% of all viral phyla compared to dsRNA-based methods. dsRNA-based methods were able to detect not only RNA viruses, but also DNA viruses except those from *Prepasmiviricota* phylum. In addition, the dsRNA-based method captured dsDNA-RT viruses (*Artverviricota* phylum) and non-enveloped ssDNA (*Phixiviricota* phylum), that VANA method missed. This result was supported by Trubl *et al.*^5^ who refined the VANA extraction-based approach to better capture both ssDNA and dsDNA viruses^5^. Overall, these results were also reinforced through the beta diversity analysis, which indicated that dsRNA-based methods (CCC-dsRNA and RT-dsRNA) and VANA method captured distinct microbial communities.

In both soil types, the dsRNA-based methods yielded an average between 5000 - 26000 viral contigs per sample. In comparison, the only soil RNA-viromics study using deep sequencing to capture soil RNA viruses identified 3462 viral contigs^10^. The primary dominant phyla in our dsRNA-based methods were *Pisuviricota* and *Duplornaviricota*. This result differs from the soil RNA-viromics results, which identified a few number of dsRNA viruses of *Duplornaviricota* phylum^10^. Also, according a recent study that compiled several datasets, *Lenarviricota* and *Pisuviricota* phylum were prevalent in soils^14^. A possible explanation of this unexpected result may be that dsRNA method favor virus with dsRNA genome whereas metranscriptomic method favors virus with single-stranded RNA genome. Additionally, the denaturing step in the dsRNA method during cDNA synthesis may facilitate the reverse transcription of those sequences. This discrepancy need to be addressed though comparative study using mock sample sets. Contrary to VANA method, dsRNA-based methods detected viruses of *Partitiviridae* family in both soil types. *Partitiviridae* are bisegmented dsRNA viruses that can infect plants, fungi, and protozoa. These viruses only spread intracellularly during spore formation in fungi, which would explain why VANA method was not able to captured them^49^. In addition, the dsRNA method captured virus of *Retroviridae* taxon, a family of RNA viruses that possess a reverse transcriptase enzyme to produce DNA from RNA genome. To our knowledge, this virus family has never being reported in agriculture soil in the published literature even though viruses of this family are known to be widespread in nature^50^. The only reporting of this family in agricultural soil comes from a recent global study that used a novel bioinformatics approach within the new version of IMG/VR platform to search viruses in existing metaviromics datasets^51^.

Indeed, the use of metaviromics and high performance computing has increased our understanding of soil RNA virus diversity in five of the six main phyla of the *Riboviria* realm^11, 18^. Using dsRNA-based methods and RdRp-based phylogenetic analysis, an expanded fine-scale RNA viral diversity was revealed, with the six phyla containing RNA viruses within *Riboviria* viral realm detected compared to metaviromics studies^10, 11, 19^. We used two complementary approaches for detection of novel RdRps within the *Riboviria* phyla. We detected a total of 3814 RdRps whereas the two core soil metaviromics studies focusing on RdRps^10, 11^ detected 3422 and 3471 RdRps, respectively. Within our detected RdRp sequences, 2944 were unique according to the annotation of functional RdRps protein motifs derived from the recently published RdRp-Scan workflow and database^52^. Among these unique RdRps, 284 were putatively novels according to the multi sequences alignment analysis of the conserved A, B and C protein motifs of the RdRps domains.

These novels RdRps fell into *Duplornaviricota* phylum, *Pisuviricota*, *Kitrinoviricota*, *Lenarviricota* and undefined phyla. The two novel viral RdRps from the undefined phyla (*incertae sedis*) belong to *Birnaviridae* family, which, to our knowledge, have never been reported in agricultural soil before. *Birnaviridae* viruses are non-enveloped bi-segmented double-stranded (dsRNA) within *Riboviria* realm and do not employ a core particle in their replication cycle^53^, thus would unlikely be captured through the recently published RNA-viromics, metaviromics or VANA methods^10, 11, 18, 19^. Interestingly, we found these viruses in both soil types (organic and mineral), likely due to their host abundance, as the four genera of the *Birnaviridae* family infect insects, fishes, and birds^54^.

Furthermore, with dsRNA-based method we identified 37 viral contigs (with six unique RdRps) from the negative-sense RNA (-ssRNA) *Negarnaviricota* phylum, compared to only four viral contigs identified in the RNA-viromics study^10^. These -ssRNA viruses are primarily lipid-enveloped, and may lack nucleocapsid proteins, making them difficult to detect using RNA-viromics approaches^55^.

In conclusion, this study examined the effectiveness of two nucleic acid extraction methods, dsRNA and VANA, in characterizing the soil virome. The results indicated that the dsRNA method was more effective than the VANA approach in capturing the soil viral diversity. Moreover, the VANA and metaviromics methods are limited to detecting either DNA or RNA viruses, while the dsRNA method has proven to detect both DNA and RNA viruses. The study emphasizes the importance of selecting an appropriate extraction method based on the purpose of a given study. For researchers looking to identify DNA viruses, the VANA method may be suitable even though it missed non-enveloped ssDNA and dsDNA viruses, while those seeking RNA and DNA viruses should consider using the dsRNA method. Even though dsRNA-based method lacked of detecting DNA viruses of *Preplasmiviricota* phylum, it offers a better profile and coverage of the soil virome compared to existing methods (VANA and metaviromics). By breaking out the limitation associated with cost-effectiveness of using metaviromics to study soil RNA viruses, this work opens the door for intensive screening of soil RNA viruses diversity and ecology. It is a cost-effective method for massive screening of soil viruses and a proven tool to expand RNA viruses diversity. Furthermore, compare to existing soil-viromics methods, sequencing dsRNA can give us an indication on virus that actively replicating in soil, because dsRNA is mainly produced as an intermediate during the viral replication process, except for viruses that has dsRNA as primary genome. The ds-RNA-based method can have broader impact beyond the soil environment and be adapted for in human gut or aquatic environments to characterize RNA viruses, which according to a recent studies are significantly less studied mostly because of their lack of stability and the difficulty of their identification in metaviromics sequencing^13, 56, 57^. Moreover, this study highlights the known need of novel computational analysis methods for soil virus diversity studies and has developed the SOVAP workflow, which is a freely available open-access tool that can be adapted to study virus diversity and ecology.

Further steps would need a close comparison between the dsRNA-based method and RNA-based method (metaviromics approach) using the same samples to determine which method is preferable based on the targeted RNA virus groups and if combining both methods may provide a more comprehensive virome characterization. The developed dsRNA-based methods can be tested using nanopore sequencing to make more affordable.

## Methods

### Sample collection

The samples used for this study were collected from two different experimental farms Agriculture and Agri-Food Canada’s in Quebec, Canada. The first farm is a vineyard in mineral soil located in Frelighsburg and the four soil samples were collected at 5-15cm depths. The second farm is in organic soil in Sainte-Clotilde and the four samples were collected at 0-10 cm. The samples were properly stored at −20 °C until proceeding with nucleic acids extraction. Each of the eight samples was homogenized and divided in the three different groups corresponding to the three extraction methods, VANA, CCC-dsRNA and RPT-dsRNA, described in the following sections. The soil samples were submitted to AgroEnviroLab for analysis of the soil’s pH level and measurement of key exchangeable minerals, such as aluminum (Al), copper (Cu), zinc (Zn), manganese (Mn), and iron (Fe) (Supplementary Table 1).

### dsRNA extraction method using RNeasy Power Soil kit (RPT-dsRNA extraction)

In this study, total RNA was extracted from the 8 soil samples, each containing 2 g, using the RNeasy Power Soil Total RNA kit (Qiagen, Hilden, Germany) as per the manufacturer’s instructions. The resulting pellet was resuspended in 100 μl of RNase-free water. Enzymatic digestion was then carried out for all 8 samples using DNase I (Thermo scientific) at a final concentration of 10 U/μl and RNase T1 (Thermo scientific) at a final concentration of 10 U/μl for 30 minutes at 37°C. The purified dsRNA was obtained using the RNA mini elute kit (Qiagen, Hilden, Germany). The dsRNA was denatured by incubating it at 99 °C for 5 minutes, followed by immediate cooling on ice for 2 minutes. cDNA was then generated by incubating the purified dsRNA with random hexamers using the SuperScript IV cDNA Synthesis kit (Thermo scientific), followed by bead purification. Finally, the resulting cDNA was quantified using the Qubit dsDNA high-sensitivity kit (Fig. 8).

**Figure 8:**
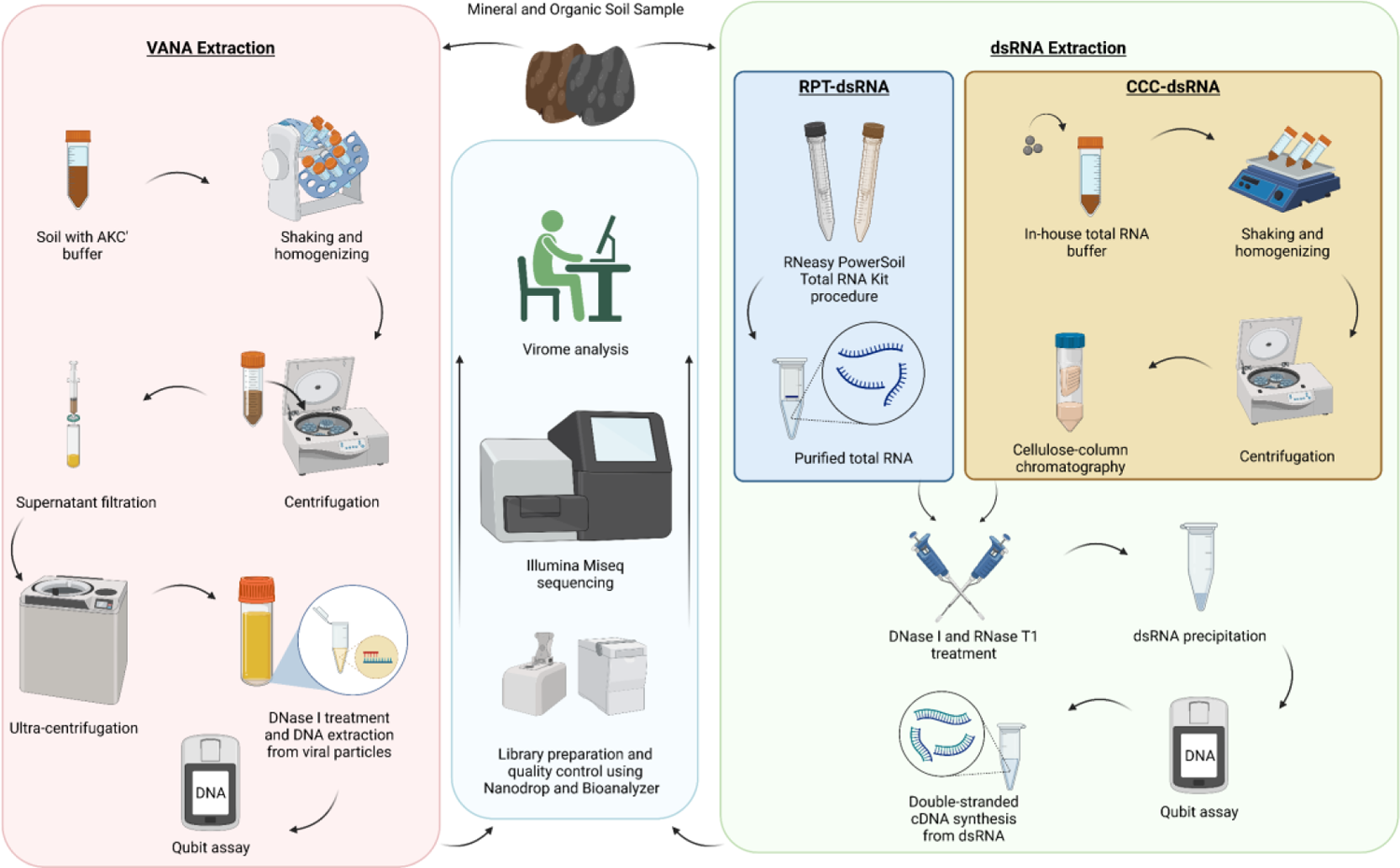
Overview of the workflow of the three extraction methods from two soil types (mineral and organic). Three extraction methods were used to extract virus nucleic acids from the soil samples, which included two dsRNA extraction methods: the RNeasy Power Soil Total RNA kit (RPT-dsRNA) and Cellulose-Column Chromatography (CCC-dsRNA), and the Virion Associated Nucleic Acid approach (VANA) to extract viral DNA. After nucleic acid extraction and quantification, samples were prepared for library construction and quality control using Bioanalyzer, followed by sequencing on the Illumina MiSeq platform.

### dsRNA extraction using cellulose-column chromatography (CCC-dsRNA extraction)

To isolate dsRNA using the cellulose-column chromatography method, total RNA was extracted from 2 and 1.5 grams of mineral and organic soil, respectively, using our optimized extraction protocol^58^. The extracted RNA was then subjected to a modified version of previously reported methods to isolate dsRNA^59–61^. Supernatant containing total RNA was mixed with absolute ethanol to obtain a solution of 16% ethanol. Furthermore, cellulose slurry was prepared using optimized chromatography buffer (50 mM Tris base, 100 mM NaCl, 16% ethanol, pH ∼ 11) at a ratio of 200 mg cellulose per ml and mixed for 10 minutes to equilibrate the cellulose powder (Sigmacell Cellulose, Type 101). 700 µl of the slurry was loaded into the PES centrifugal filter column (Sartorius, 0.8 µm) and centrifuged at 10,000 g for 10 seconds. Following that, the supernatant containing 16% ethanol was loaded on the column and centrifuged again. To further minimize humic contamination, four washes were performed using chromatography buffer to remove the majority of humic compounds from the column. To remove residual ethanol, the column was centrifuged at the same speed for 15 seconds prior to elution. The column was eluted with chromatography buffer (without ethanol, pH 8) followed by precipitation with 5 M LiCl final concentration and one volume of isopropanol for two hours. Upon dissolving the pellet in 20 µl ultrapure water and prior to cDNA synthesis, if traces of humic acid were detected, a final purification was conducted using the OneStep PCR Inhibitor Removal Kit (Zymo Research). After denaturing the dsRNA the cDNA was then generated as described in the previous section (RPT-dsRNA extraction). The resulting cDNA was then quantified using the Qubit dsDNA high-sensitivity kit.

### Virion associated Nucleic Acid extraction (VANA)

To extract virion-associated nucleic acids, we used the protocol described in Santos-Medellin *et al.*^62^. Briefly, for each 50 g of soil sample, two 50 mL conical tubes were filled with 25 g of soil and 37.5 mL of 0.22 µm filtered AKC’ extraction buffer. The resulting mixture was homogenized, shaken on an orbital shaker at 400 RPM for 15 minutes, vortexed for 3 minutes, and centrifuged at 4,700 x g for 15 minutes. The supernatant was filtered through a 0.22 µm filter. The tube was then centrifuged at 32,000 x g for 3 hours at 4°C, and the resulting pellets containing the viral fraction were resuspended in 200 µl of ultrapure water. To remove free DNA, the resuspended pellets were treated with 30 units of DNase I (Thermo scientific) and 30 µl of 10X DNase buffer for 2 hours at room temperature, and the reaction was quenched with DNase stop solution. Nucleic acid was extracted using the Dneasy Power Soil Pro kit (Qiagen). The extracted DNA was quantified by employing a Qubit spectrofluorometry with Qubit dsDNA High Sensitivity kit. (Fig. 8).

### Library construction and sequencing

The libraries were prepared using the Nextera DNA library prep kit and 1 ng of input double-stranded cDNA. Subsequently, the Illumina Miseq platform sequencer was used to perform sequencing with the MiSeq Reagent Kits v3.

#### Virome data analysis

Adapters and low-quality reads were removed from raw sequencing data using Fastp v0.23.2^63^. Subsequently, the resulting high-quality reads were decontaminated using Centrifuge v1.0.4^64^ with a custom-built database. The clean reads were then assembled into contigs using MEGAHIT v1.2.9^65^. The recently introduced geNomad v1.3.3 tool was utilized for the identification of viral contigs^66^, which were then clustered using CD-HIT v4.8.1^67, 68^ for further analysis. The abundance of the viral contigs was estimated using BWA v0.7.17-r1188^69^, and an in-house script was used to calculate abundance values for each contig. The clustered viral contigs were annotated using DIAMOND (blastx) v2.1.4^70^ with the in-house indexed IMG/VR v4 database^71^. The resulting output was visualized using R v4.2.0^72^, and the taxonomy was plotted with Tidyverse v1.3.2 sub-packages^73^ for each taxonomy rank^74^.

The alpha-diversity analysis, including species richness, Shannon, and Simpson indices, was calculated and visualized using the Vegan v2.6-4^75^ and Tidyverse packages, respectively. Detailed steps and codes for the entire process are available in the GitHub repository (https://github.com/poursalavati/SOVAP_Soil_dsRNA)^41^. Additionally, a similar analysis was conducted by concatenating all biological replicates of each method and analyzing them together, following the same methodology as described earlier. Moreover, beta-diversity analysis was conducted to evaluate the dissimilarity and similarity of viral communities at each taxonomy rank between the viromics methods using the Bray-Curtis distance matrix, and principal coordinate analysis (PCoA) was employed to visualize the results^74^. Permutational multivariate analysis of variance (PERMANOVA) was used to determine the statistical significance of the observed differences.

#### Phylogenetic analysis

Diamond blastp and RdRp-Scan datasets^27^ were used to search for RdRp genes in the identified viral proteins from geNomad (> 70 amino acids). A second analysis of the resulting hits was conducted using HMMSearch v3.3.2^76^ with the RdRp-Scan Hidden Markov model (HMM) profile database to improve the accuracy of putative RdRp sequences. Then contigs with E values < 0.001 and scores >50 were clustered with CD-Hit to remove redundant sequences (with 95% average amino acid identity). The putative candidates were then categorized into different taxa and aligned using MAFFT v7.515^77^ and Clustal Omega v1.2.4^78^ to the RdRp-Scan pre-built backbone trees, as well as previously reported RdRp sequences from the Wolf^79^, Starr^20^, and Hillary^80^ studies (WSH) for each individual taxa, respectively. Trees were generated with FastTree (MP, Multi-threaded) v2.1.11^81^ and visualized with iToL v6.7.3^82^. We used a multi-step approach to re-analyze RdRp candidates and identify novel sequences. First, we identified three key RdRp motifs (A, B, and C) using a motif database^27^ and CLC Genomics Workbench v22 (QIAGEN). Next, we performed multiple sequence alignment (MSA) with MegAlign Pro v17.4.1 (DNAStar Lasergene software) to identify conserved regions and potential variations among the candidates of each *Riboviria* taxa. Finally, we visualized the MSA results and conserved motifs logos using Inkscape v1.21 (https://inkscape.org). This approach allowed us to effectively analyze the candidates and identify important functional motifs in potential novel instances.

## Data availability

The sequencing data supporting this research have been deposited in the NCBI Sequence Read Archive (SRA) repository under BioProject accession number PRJNA948674. Detailed bioinformatics workflow (SOVAP scripts), along with the generated databases, are available at https://github.com/poursalavati/SOVAP, https://zenodo.org/record/7758200 and https://zenodo.org/record/7747335, respectively. Custom R scripts for downstream data analysis, along with results of abundance, alignments, phylogenetic trees, and identified RdRp sequences from all datasets are accessible on the following GitHub repository: https://github.com/poursalavati/SOVAP_Soil_dsRNA.

## Acknowledgments

The authors gratefully acknowledge the administrative support from the CRD of Saint-Jean-sur-Richelieu team, including Vicky Toussaint and Mélanie Cadieux teams. We would also like to express our gratitude to Joel Lafond-Lapalme for his valuable support in maintaining the bioinformatics system. Furthermore, we acknowledge Eric Courchesne and his team for their support and management of the experimental vineyard.

## Funding

This research was funded by Agriculture and Agri-Food Canada under the Ensuring water and soil resources sustainability (J-002302) and ecosystem productivity and resiliency (J-001792) programs.

## Notes

https://github.com/poursalavati/SOVAP

https://zenodo.org/record/7758200

https://zenodo.org/record/7747335

https://github.com/poursalavati/SOVAP_Soil_dsRNA

### Competing Interest Statement

The authors have declared no competing interest.

## References

1. Edwards RA, Rohwer F. Viral metagenomics. Nature Reviews Microbiology 3, 504–510 (2005).

2. Williamson KE, Fuhrmann JJ, Wommack KE, Radosevich M. Viruses in Soil Ecosystems: An Unknown Quantity Within an Unexplored Territory. Annual Review of Virology 4, 201–219 (2017).

3. Emerson JB, et al. Host-linked soil viral ecology along a permafrost thaw gradient. Nature Microbiology 3, 870–880 (2018).

4. Pratama AA, Elsas JDv. The ‘Neglected’ Soil Virome – Potential Role and Impact. Trends in Microbiology 26, 649–662 (2018).

5. Trubl G, et al. Towards optimized viral metagenomes for double-stranded and single-stranded DNA viruses from challenging soils. PeerJ 7, e7265 (2019).

6. Williamson KE, Radosevich M, Wommack KE. Abundance and diversity of viruses in six Delaware soils. Appl Environ Microbiol 71, 3119–3125 (2005).

7. Ackermann HW. 5500 Phages examined in the electron microscope. Arch Virol 152, 227–243 (2007).

8. Reavy B, et al. Distinct circular single-stranded DNA viruses exist in different soil types. Appl Environ Microbiol 81, 3934–3945 (2015).

9. Liang X, et al. Viral abundance and diversity vary with depth in a southeastern United States agricultural ultisol. Soil Biology and Biochemistry 137, 107546 (2019).

10. Hillary LS, Adriaenssens EM, Jones DL, McDonald JE. RNA-viromics reveals diverse communities of soil RNA viruses with the potential to affect grassland ecosystems across multiple trophic levels. ISME Commun 2, 34 (2022).

11. Starr EP, Nuccio EE, Pett-Ridge J, Banfield JF, Firestone MK. Metatranscriptomic reconstruction reveals RNA viruses with the potential to shape carbon cycling in soil. Proc Natl Acad Sci U S A 116, 25900–25908 (2019).

12. Koonin EV, Krupovic M, Dolja VV. The global virome: How much diversity and how many independent origins? Environ Microbiol 25, 40–44 (2023).

13. Cobbin JC, Charon J, Harvey E, Holmes EC, Mahar JE. Current challenges to virus discovery by meta-transcriptomics. Curr Opin Virol 51, 48–55 (2021).

14. Jansson JK, Wu R. Soil viral diversity, ecology and climate change. Nature Reviews Microbiology, (2022).

15. Andika IB, Kondo H, Sun L. Interplays between Soil-Borne Plant Viruses and RNA Silencing- Mediated Antiviral Defense in Roots. Frontiers in Microbiology 7, (2016).

16. Zhang R, Liu S, Chiba S, Kondo H, Kanematsu S, Suzuki N. A novel single-stranded RNA virus isolated from a phytopathogenic filamentous fungus, Rosellinia necatrix, with similarity to hypo-like viruses. Frontiers in Microbiology 5, (2014).

17. Chen YM, et al. RNA viromes from terrestrial sites across China expand environmental viral diversity. Nat Microbiol 7, 1312–1323 (2022).

18. Wu R, et al. Moisture modulates soil reservoirs of active DNA and RNA viruses. Commun Biol 4, 992 (2021).

19. Wolf YI, et al. Origins and Evolution of the Global RNA Virome. mBio 9, (2018).

20. Starr EP, Nuccio EE, Pett-Ridge J, Banfield JF, Firestone MK. Metatranscriptomic reconstruction reveals RNA viruses with the potential to shape carbon cycling in soil. Proceedings of the National Academy of Sciences 116, 25900–25908 (2019).

21. Shi M, et al. Redefining the invertebrate RNA virosphere. Nature 540, 539–543 (2016).

22. Callanan J, Stockdale SR, Shkoporov A, Draper LA, Ross RP, Hill C. Expansion of known ssRNA phage genomes: From tens to over a thousand. Science Advances 6, eaay5981 (2020).

23. Wolf YI, et al. Doubling of the known set of RNA viruses by metagenomic analysis of an aquatic virome. Nature Microbiology 5, 1262–1270 (2020).

24. Neri U, et al. Expansion of the global RNA virome reveals diverse clades of bacteriophages. Cell 185, 4023–4037.e4018 (2022).

25. Koonin EV, et al. Global Organization and Proposed Megataxonomy of the Virus World. Microbiology and molecular biology reviews: MMBR 84, e00061–00019 (2020).

26. Koonin EV. The phylogeny of RNA-dependent RNA polymerases of positive-strand RNA viruses. Journal of General Virology 72, 2197–2206 (1991).

27. Charon J, Buchmann JP, Sadiq S, Holmes EC. RdRp-scan: A bioinformatic resource to identify and annotate divergent RNA viruses in metagenomic sequence data. Virus Evolution 8, veac082 (2022).

28. Sakaguchi S, et al. NeoRdRp: A Comprehensive Dataset for Identifying RNA-dependent RNA Polymerases of Various RNA Viruses from Metatranscriptomic Data. Microbes and Environments 37, ME22001 (2022).

29. Edgar RC. Muscle5: High-accuracy alignment ensembles enable unbiased assessments of sequence homology and phylogeny. Nature Communications 13, 6968 (2022).

30. Ferrero DS, Falqui M, Verdaguer N. Snapshots of a Non-Canonical RdRP in Action. Viruses 13, 1260 (2021).

31. Sutela S, Poimala A, Vainio EJ. Viruses of fungi and oomycetes in the soil environment. FEMS Microbiol Ecol 95, (2019).

32. Kumar M, Carmichael GG. Antisense RNA: function and fate of duplex RNA in cells of higher eukaryotes. Microbiol Mol Biol Rev 62, 1415–1434 (1998).

33. Weber F, Wagner V, Rasmussen SB, Hartmann R, Paludan SR. Double-stranded RNA is produced by positive-strand RNA viruses and DNA viruses but not in detectable amounts by negative-strand RNA viruses. J Virol 80, 5059–5064 (2006).

34. Chen YG, Hur S. Cellular origins of dsRNA, their recognition and consequences. Nat Rev Mol Cell Biol, 286–301 (2022).

35. Tzanetakis IE, Martin RR. Fragaria chiloensis cryptic virus: A New Strawberry Virus Found in Fragaria chiloensis Plants from Chile. Plant Disease 89, 1241–1241 (2005).

36. Tzanetakis IE, Martin RR. A new method for extraction of double-stranded RNA from plants. Journal of Virological Methods 149, 167–170 (2008).

37. Gaafar YZA, Ziebell H. Comparative study on three viral enrichment approaches based on RNA extraction for plant virus/viroid detection using high-throughput sequencing. PLoS One 15, e0237951 (2020).

38. Tzanetakis IE, Martin RR. A new method for extraction of double-stranded RNA from plants. J Virol Methods 149, 167–170 (2008).

39. Fall ML, Xu D, Lemoyne P, Moussa IEB, Beaulieu C, Carisse O. A Diverse Virome of Leafroll- Infected Grapevine Unveiled by dsRNA Sequencing. Viruses 12, (2020).

40. Blouin AG, Ross HA, Hobson-Peters J, O’Brien CA, Warren B, MacDiarmid R. A new virus discovered by immunocapture of double-stranded RNA, a rapid method for virus enrichment in metagenomic studies. Mol Ecol Resour 16, 1255–1263 (2016).

41. Poursalavati A. SOVAP v.1.3: Soil Virome Analysis Pipeline.). 1.3 edn. Zenodo (2023).

42. Tange O. GNU Parallel 20221122 (’XepcóH’).). Zenodo (2022).

43. Hillary LS, Adriaenssens EM, Jones DL, McDonald JE. RNA-viromics reveals diverse communities of soil RNA viruses with the potential to affect grassland ecosystems across multiple trophic levels.) (2022).

44. Jansson JK, Wu R. Soil viral diversity, ecology and climate change. Nat Rev Microbiol, (2022).

45. Schulz F, Abergel C, Woyke T. Giant virus biology and diversity in the era of genome- resolved metagenomics. Nat Rev Microbiol 20, 721–736 (2022).

46. Mushegian AR. Are There 10(31) Virus Particles on Earth, or More, or Fewer? J Bacteriol 202, (2020).

47. Walker PJ, et al. Recent changes to virus taxonomy ratified by the International Committee on Taxonomy of Viruses (2022). Arch Virol 167, 2429–2440 (2022).

48. Steward GF, Culley AI, Mueller JA, Wood-Charlson EM, Belcaid M, Poisson G. Are we missing half of the viruses in the ocean? The ISME Journal 7, 672–679 (2013).

49. Vainio EJ, et al. ICTV Virus Taxonomy Profile: Partitiviridae. J Gen Virol 99, 17–18 (2018).

50. Coffin J, et al. ICTV Virus Taxonomy Profile: Retroviridae 2021. J Gen Virol 102, (2021).

51. Camargo AP, et al. IMG/VR v4: an expanded database of uncultivated virus genomes within a framework of extensive functional, taxonomic, and ecological metadata. Nucleic Acids Res 51, D733–D743 (2023).

52. Charon J, Buchmann JP, Sadiq S, Holmes EC. RdRp-scan: A bioinformatic resource to identify and annotate divergent RNA viruses in metagenomic sequence data. Virus Evol 8, veac082 (2022).

53. Dalton RM, Rodríguez JF. Rescue of Infectious Birnavirus from Recombinant Ribonucleoprotein Complexes. PLoS ONE 9, e87790 (2014).

54. Delmas B, et al. ICTV virus taxonomy profile: Birnaviridae. In: Microbiology Society) (2019).

55. Compans RW, et al. Escaping from the Cell: Assembly and Budding of Negative-Strand RNA Viruses. In: Biology of Negative Strand RNA Viruses: The Power of Reverse Genetics (ed Kawaoka Y). Springer Berlin Heidelberg (2004).

56. Callanan J, Stockdale SR, Shkoporov A, Draper LA, Ross RP, Hill C. Expansion of known ssRNA phage genomes: From tens to over a thousand. Sci Adv 6, eaay5981 (2020).

57. Cao Z, Sugimura N, Burgermeister E, Ebert MP, Zuo T, Lan P. The gut virome: A new microbiome component in health and disease. EBioMedicine 81, 104113 (2022).

58. Poursalavati A, Javaran VJ, Laforest-Lapointe I, Fall M. Soil metatranscriptomics: An improved RNA extraction method toward functional analysis using nanopore direct RNA sequencing. Phytobiomes Journal, (2023).

59. Fall ML, Xu D, Lemoyne P, Moussa IEB, Beaulieu C, Carisse O. A diverse virome of leafroll- infected grapevine unveiled by dsRNA sequencing. Viruses 12, 1142 (2020).

60. Peyambari M, Roossinck MJ. Characterizing mycoviruses. In: Plant pathogenic fungi and oomycetes). Springer (2018).

61. Choi YG, Randles JW. Microgranular cellulose improves dsRNA recovery from plant nucleic acid extracts. BioTechniques 23, 610–611 (1997).

62. Santos-Medellin C, Zinke LA, Ter Horst AM, Gelardi DL, Parikh SJ, Emerson JB. Viromes outperform total metagenomes in revealing the spatiotemporal patterns of agricultural soil viral communities. The ISME Journal 15, 1956–1970 (2021).

63. Chen S, Zhou Y, Chen Y, Gu J. fastp: an ultra-fast all-in-one FASTQ preprocessor. Bioinformatics 34, i884–i890 (2018).

64. Kim D, Song L, Breitwieser FP, Salzberg SL. Centrifuge: rapid and sensitive classification of metagenomic sequences. Genome Research 26, 1721–1729 (2016).

65. Li D, Liu C-M, Luo R, Sadakane K, Lam T-W. MEGAHIT: an ultra-fast single-node solution for large and complex metagenomics assembly via succinct de Bruijn graph. Bioinformatics 31, 1674–1676 (2015).

66. Camargo AP, et al. You can move, but you can’t hide: identification of mobile genetic elements with geNomad. bioRxiv, 2023-2003 (2023).

67. Fu L, Niu B, Zhu Z, Wu S, Li W. CD-HIT: accelerated for clustering the next-generation sequencing data. Bioinformatics 28, 3150–3152 (2012).

68. Li W, Godzik A. Cd-hit: a fast program for clustering and comparing large sets of protein or nucleotide sequences. Bioinformatics 22, 1658–1659 (2006).

69. Li H. Aligning sequence reads, clone sequences and assembly contigs with BWA-MEM. arXiv preprint arXiv:13033997, (2013).

70. Buchfink B, Reuter K, Drost H-G. Sensitive protein alignments at tree-of-life scale using DIAMOND. Nature methods 18, 366–368 (2021).

71. Camargo AP, et al. IMG/VR v4: an expanded database of uncultivated virus genomes within a framework of extensive functional, taxonomic, and ecological metadata. Nucleic Acids Research 51, D733–D743 (2023).

72. Team RC. R: A language and environment for statistical computing (version 4.2. 0)[Programming language].). R Foundation for Statistical Computing. https://www.R-project.org (2022).

73. Wickham H, et al. Welcome to the Tidyverse. Journal of open source software 4, 1686 (2019).

74. Tao S, Zou H, Li J, Wei H. Landscapes of Enteric Virome Signatures in Early-Weaned Piglets. Microbiology Spectrum 10, e01698–01622 (2022).

75. Oksanen J, et al. vegan: Community Ecology Package. (2022).

76. Mistry J, Finn RD, Eddy SR, Bateman A, Punta M. Challenges in homology search: HMMER3 and convergent evolution of coiled-coil regions. Nucleic acids research 41, e121–e121 (2013).

77. Katoh K, Standley DM. MAFFT multiple sequence alignment software version 7: improvements in performance and usability. Molecular biology and evolution 30, 772–780 (2013).

78. Sievers F, et al. Fast, scalable generation of high-quality protein multiple sequence alignments using Clustal Omega. Molecular systems biology 7, 539 (2011).

79. Wolf YI, et al. Origins and evolution of the global RNA virome. MBio 9, e02329–02318 (2018).

80. Hillary LS, Adriaenssens EM, Jones DL, McDonald JE. RNA-viromics reveals diverse communities of soil RNA viruses with the potential to affect grassland ecosystems across multiple trophic levels. ISME Communications 2, 34 (2022).

81. Price MN, Dehal PS, Arkin AP. FastTree 2–approximately maximum-likelihood trees for large alignments. PloS one 5, e9490 (2010).

82. Letunic I, Bork P. Interactive Tree Of Life (iTOL) v5: an online tool for phylogenetic tree display and annotation. Nucleic acids research 49, W293–W296 (2021).

