## Supplementary Figs and Tables for "dsRNA-based viromics: A novel tool unveiled hidden soil viral diversity and richness"

Supplementary Table 1. Physicochemical characteristics of mineral and organic soil samples

| Properties | Units | Mineral soils | | | | Organic soils | | | |
|---|---|---|---|---|---|---|---|---|---|
| | | S1 | S2 | S3 | S4 | S1 | S2 | S3 | S4 |
| pH | - | 5.6 | 5.5 | 5.4 | 5.2 | 6.5 | 6.0 | 6.4 | 6.8 |
| Organic content | % | 5.4 | 5.5 | 5.4 | 5.2 | 72.3 | 73.7 | 71.7 | 67.6 |
| Organic carbon | % | 4.8 | 4.8 | 4.8 | 4.8 | 45.2 | 45.2 | 45.2 | 45.2 |
| CEC | meq/100g | 16.7 | 17.0 | 17.6 | 15.4 | 31.0 | 36.4 | 34.4 | 34.6 |
| P | kg/ha | 72 | 56 | 39 | 39 | 38 | 49 | 35 | 26 |
| K | kg/ha | 101 | 98 | 133 | 156 | 208 | 279 | 236 | 129 |
| Ca | kg/ha | 3236 | 3046 | 2697 | 1839 | 10636 | 10532 | 11069 | 12356 |
| Mg | kg/ha | 142 | 135 | 169 | 106 | 668 | 719 | 670 | 646 |
| Al | ppm | 994 | 1042 | 1143 | 1176 | 26 | 29 | 41 | 51 |
| Mn | ppm | 49.2 | 39.7 | 39.0 | 36.4 | 5.4 | 5.4 | 5.0 | 5.5 |
| Cu | ppm | 2.29 | 2.02 | 2.41 | 3.45 | 3.25 | 5.26 | 3.94 | 3.34 |
| Zn | ppm | 2.27 | 2.08 | 2.13 | 2.04 | 9.41 | 15.88 | 41.42 | 77.62 |
| B | ppm | 0.43 | 0.33 | 0.39 | 0.38 | 1.20 | 1.24 | 1.34 | 1.47 |
| Fe | ppm | 193 | 194 | 164 | 188 | 199 | 196 | 178 | 164 |

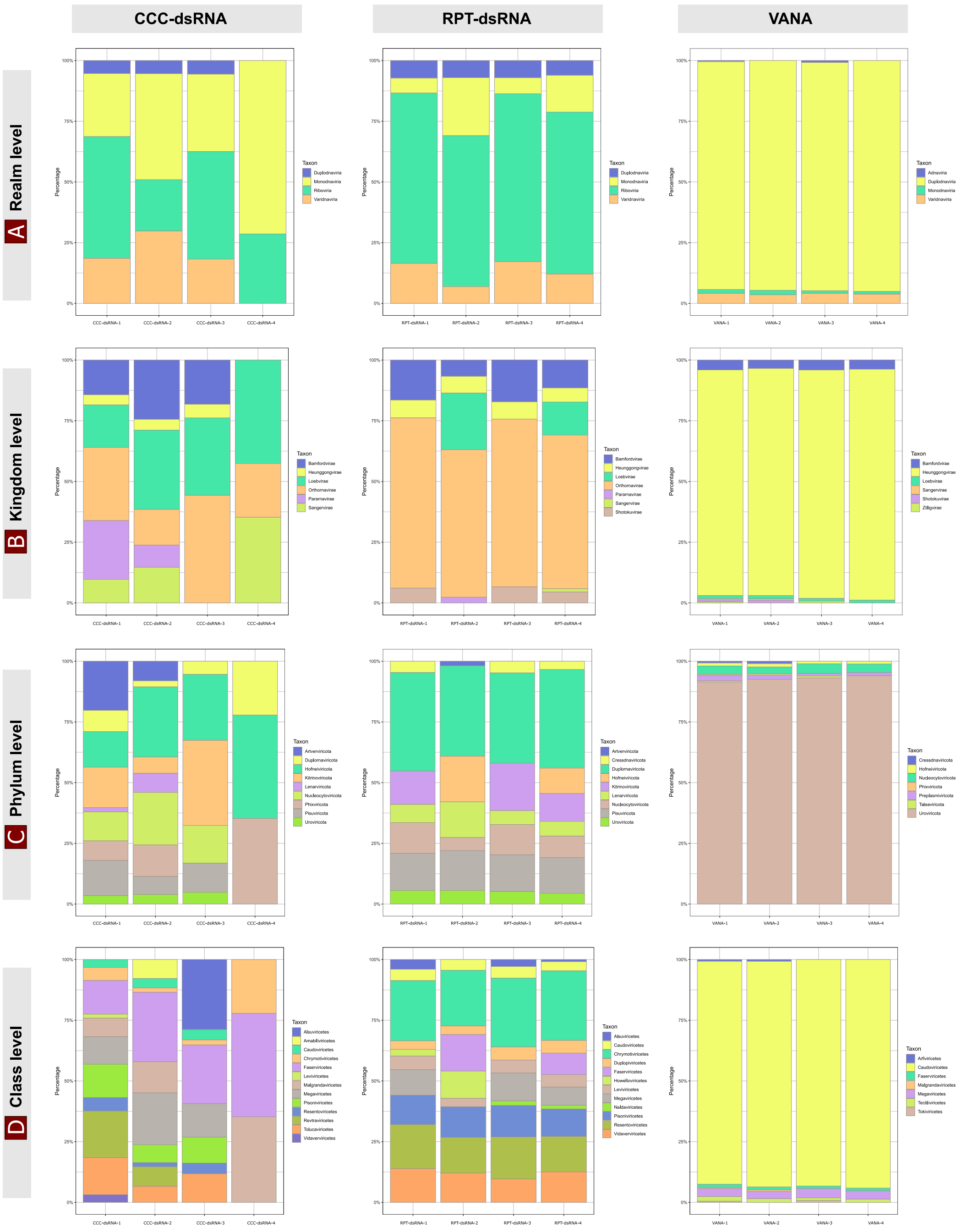


**Supplementary Figure 1: Comparison of taxonomic composition of viral communities in four organic soil sample replicates extracted using three different methods**: RNeasy Power Soil Total RNA kit (RPT-dsRNA), Cellulose-Column Chromatography (CCC-dsRNA), and the Virion Associated Nucleic Acid approach (VANA), at the taxonomic ranks of (a) realm, (b) kingdom, (c) phylum and (d) class. The stacked bar chart displays the percentage of assigned viral sequences for each taxonomic level.

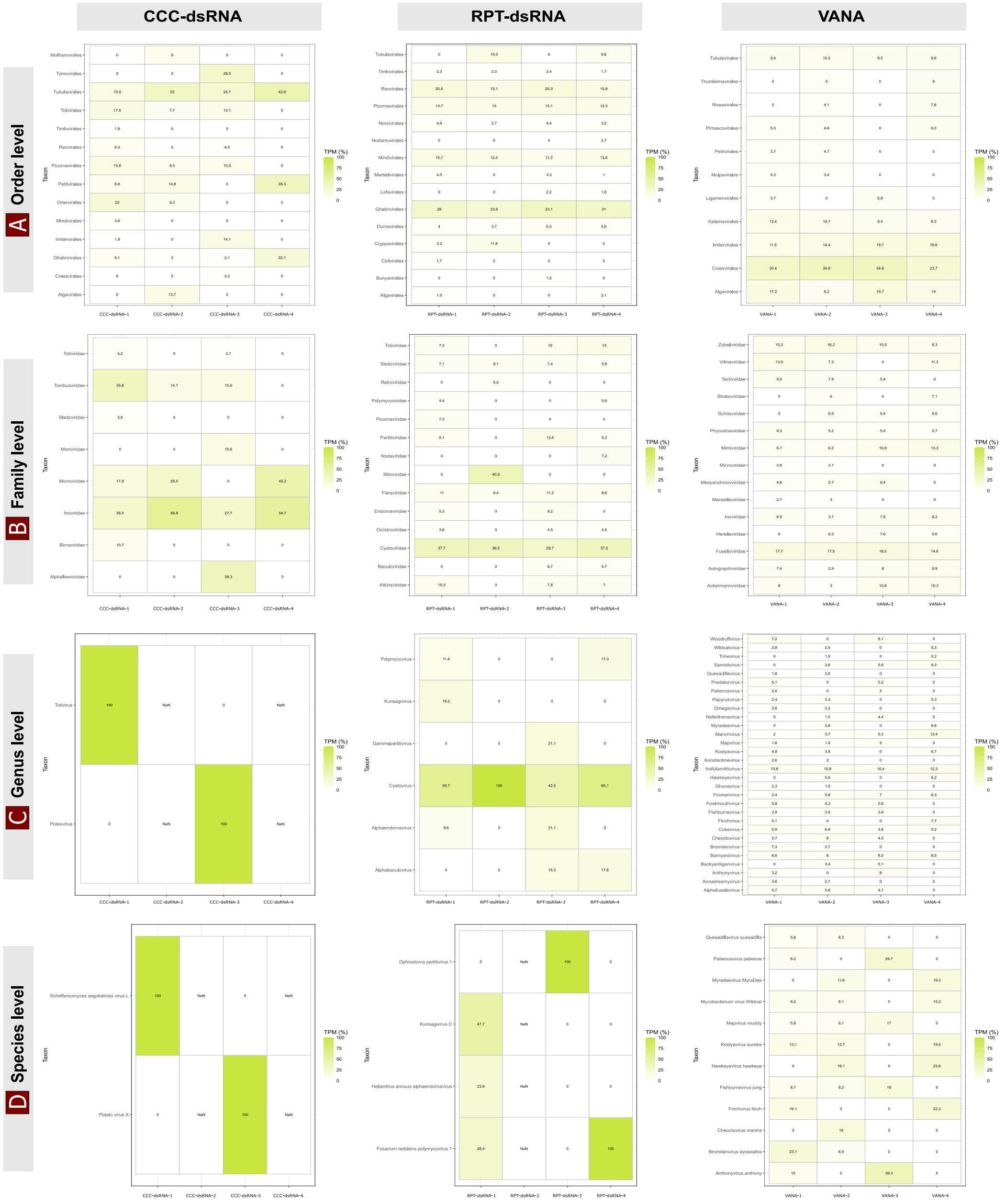


**Supplementary Figure 2: Comparison of viral taxa abundance across four organic soil sample replicates using different extraction methods**: RNeasy Power Soil Total RNA kit (RPT-dsRNA) and Cellulose-Column Chromatography (CCC-dsRNA), and the Virion Associated Nucleic Acid approach (VANA) at the (a) Order, (b) Family, (c) Genus and (d) Species levels. A heatmap shows the relative abundances determined by calculating the Transcripts per million (TPM) value of the reads counts of each viral taxa and their relative abundances in each sample replicate. The percentages of each viral taxa are indicated in the corresponding rectangles.

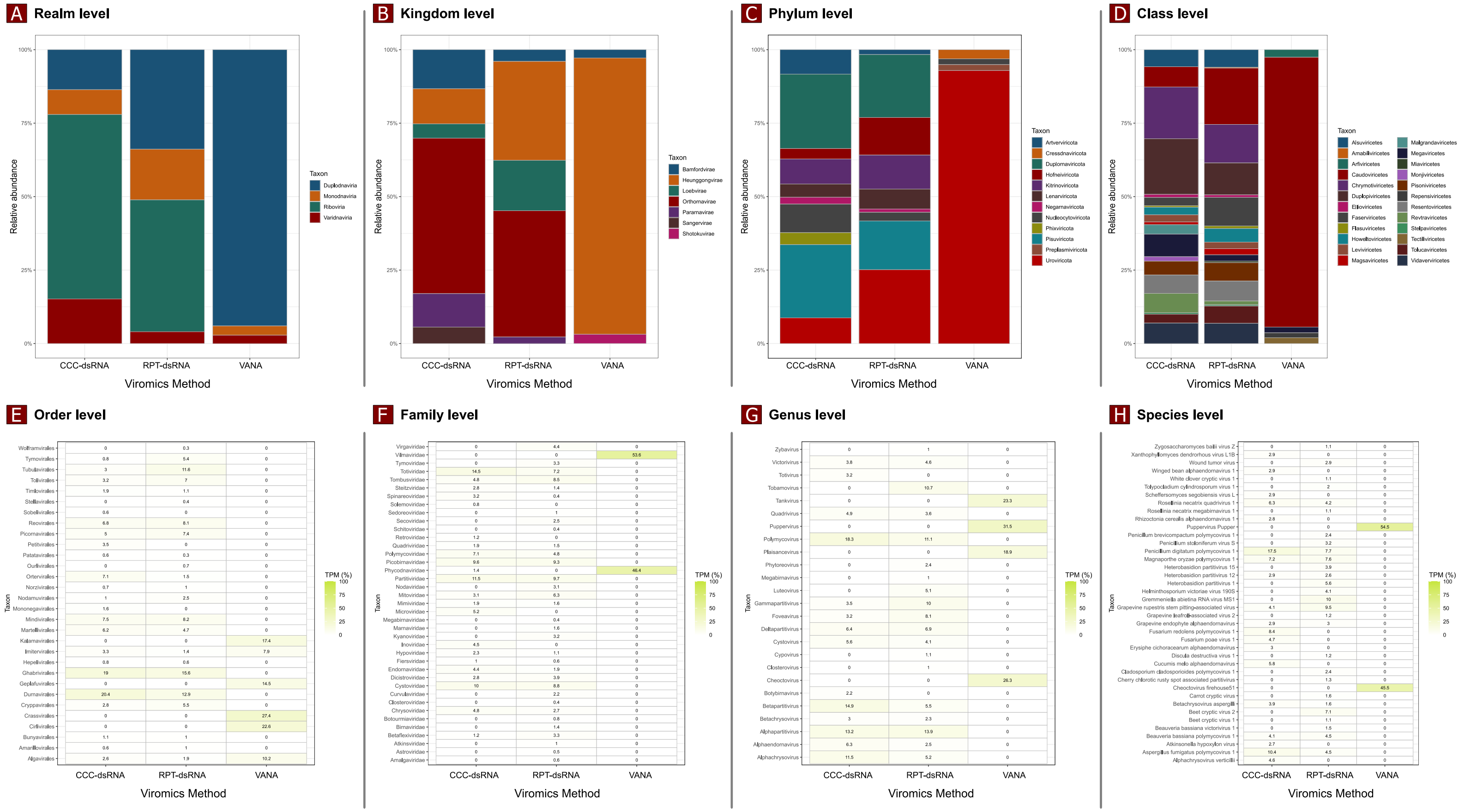


**Supplementary Figure 3: Comparison of viral taxonomic composition in combined mineral soil samples using three different extraction methods:** RNeasy Power Soil Total RNA kit (RPT-dsRNA) and Cellulose-Column Chromatography (CCC-dsRNA), and the Virion Associated Nucleic Acid approach (VANA). Taxonomic assignments are presented at various levels of classification, including (a) realm, (b) kingdom, (c) phylum, and (d) class using stacked bar charts. Heatmaps were generated to display the distribution of viral taxa at different levels, including (e) order, (f) family, (g) genus, and (h) species. The relative abundances were determined by calculating the Transcripts per million (TPM) value of the read counts for each viral taxa.

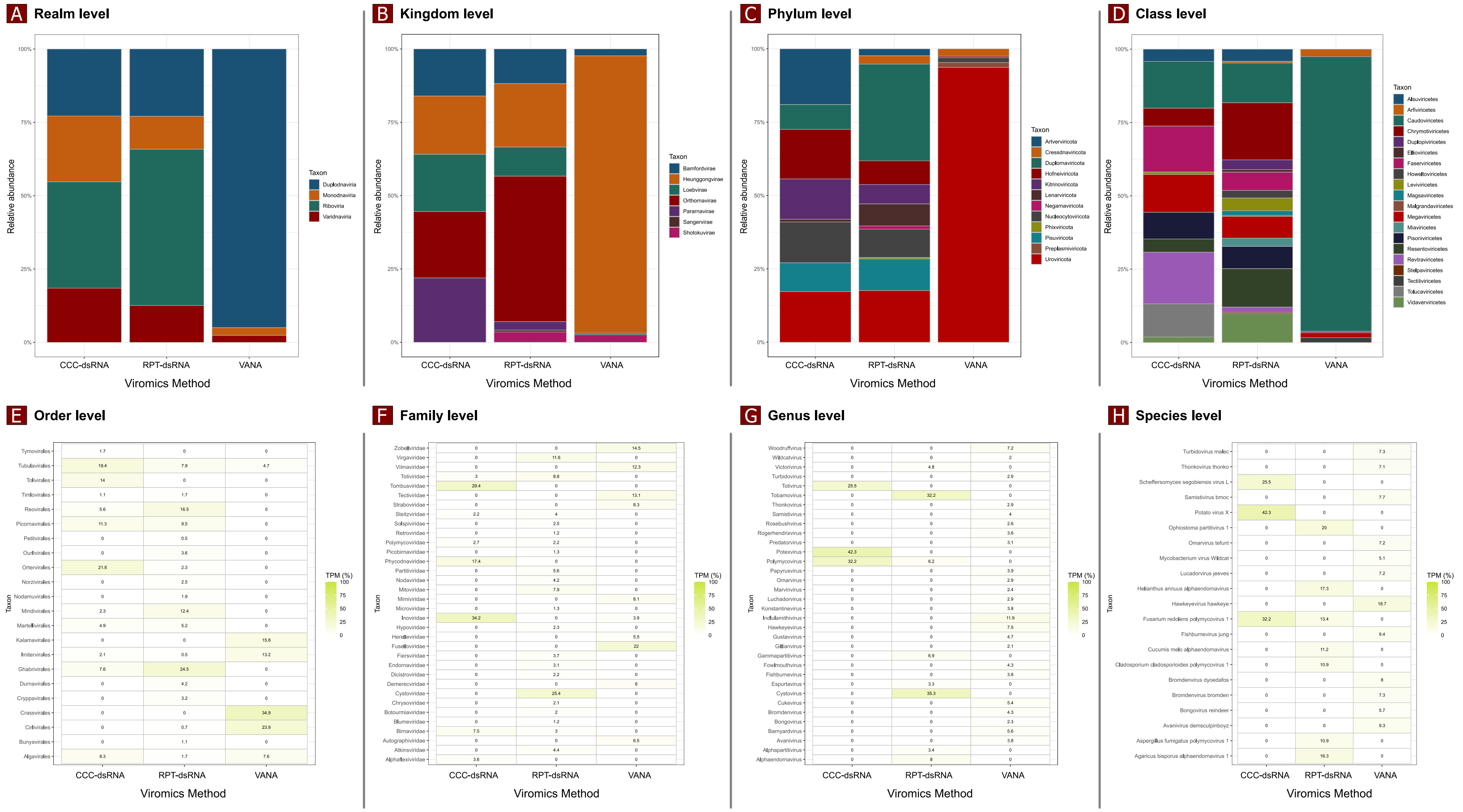


**Supplementary Figure 4: Comparison of viral taxonomic composition in combined organic soil samples using three different extraction methods:** RNeasy Power Soil Total RNA kit (RPT-dsRNA) and Cellulose-Column Chromatography (CCC-dsRNA), and the Virion Associated Nucleic Acid approach (VANA). Taxonomic assignments are presented at various levels of classification, including (a) realm, (b) kingdom, (c) phylum, and (d) class using stacked bar charts. Heatmaps were generated to display the distribution of viral taxa at different levels, including (e) order, (f) family, (g) genus, and (h) species. The relative abundances were determined by calculating the Transcripts per million (TPM) value of the read counts for each viral taxa.

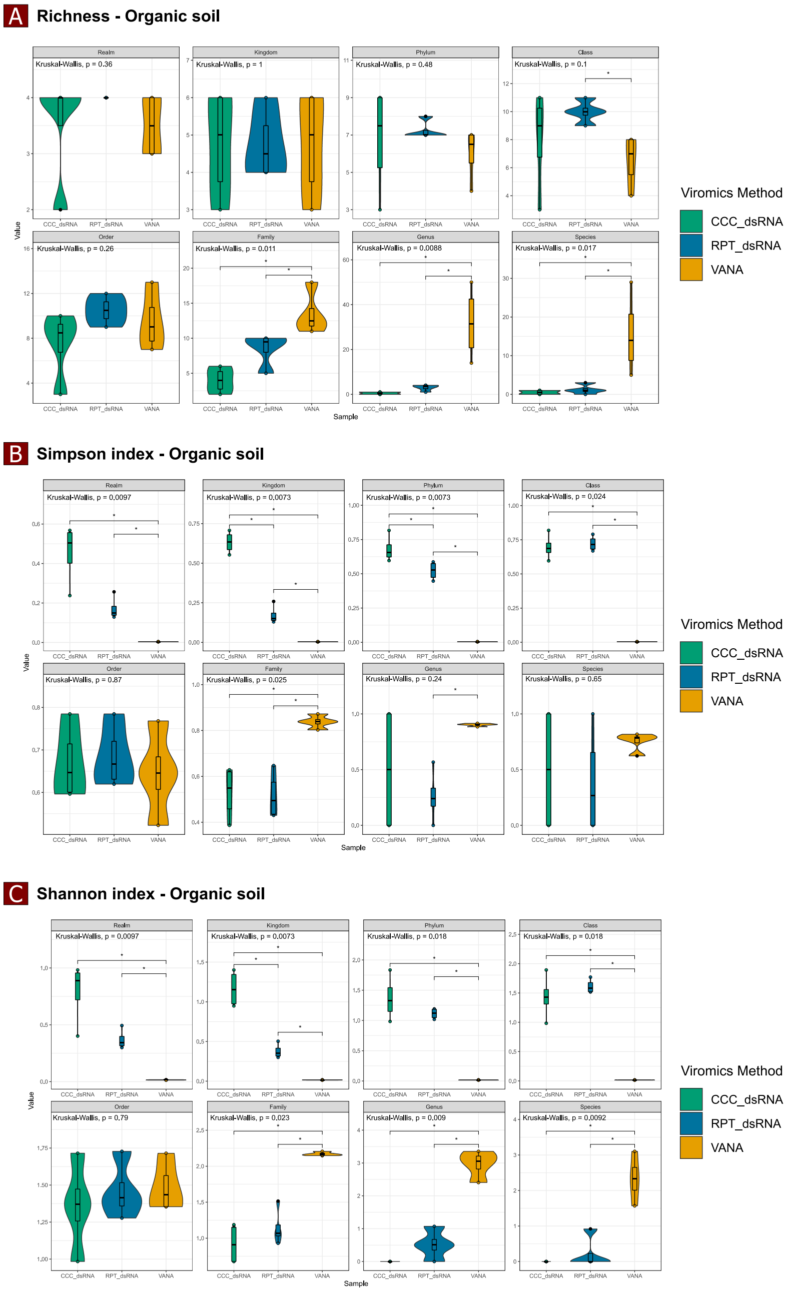


**Supplementary Figure 5: Alpha-diversity analysis of viral community composition in organic soil using three extraction methods**: RNeasy Power Soil Total RNA kit (RPT-dsRNA) and Cellulose-Column Chromatography (CCC-dsRNA), and the Virion Associated Nucleic Acid approach (VANA). Violin plots show the Richness (A), Shannon (B), and Simpson (C) indices of all biological replicates of each method, calculated using Vegan v2.6-4 and Tidyverse packages. The analysis was performed at the realm, kingdom, phylum, class, order, family, genus, and species levels.

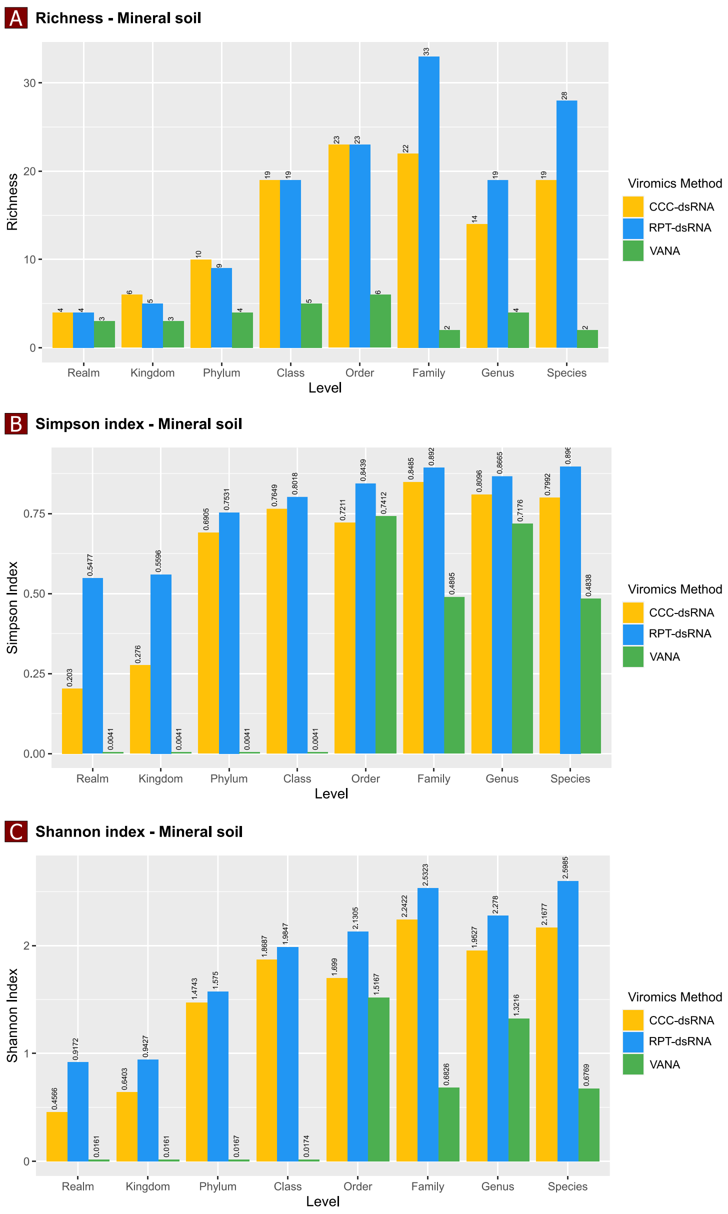


**Supplementary Figure 6: Comparison of alpha-diversity metrics for viral communities in mineral soil using three different extraction methods:** RNeasy Power Soil Total RNA kit (RPT-dsRNA), Cellulose-Column Chromatography (CCC-dsRNA), and Virion Associated Nucleic Acid (VANA). The bar chart shows richness (A), Shannon diversity (B), and Simpson diversity (C) indices for each method, calculated using Vegan v2.6-4 and Tidyverse packages. Analysis was performed at the realm, kingdom, phylum, class, order, family, genus, and species levels. The corresponding colors in the legend indicate the different extraction methods used.

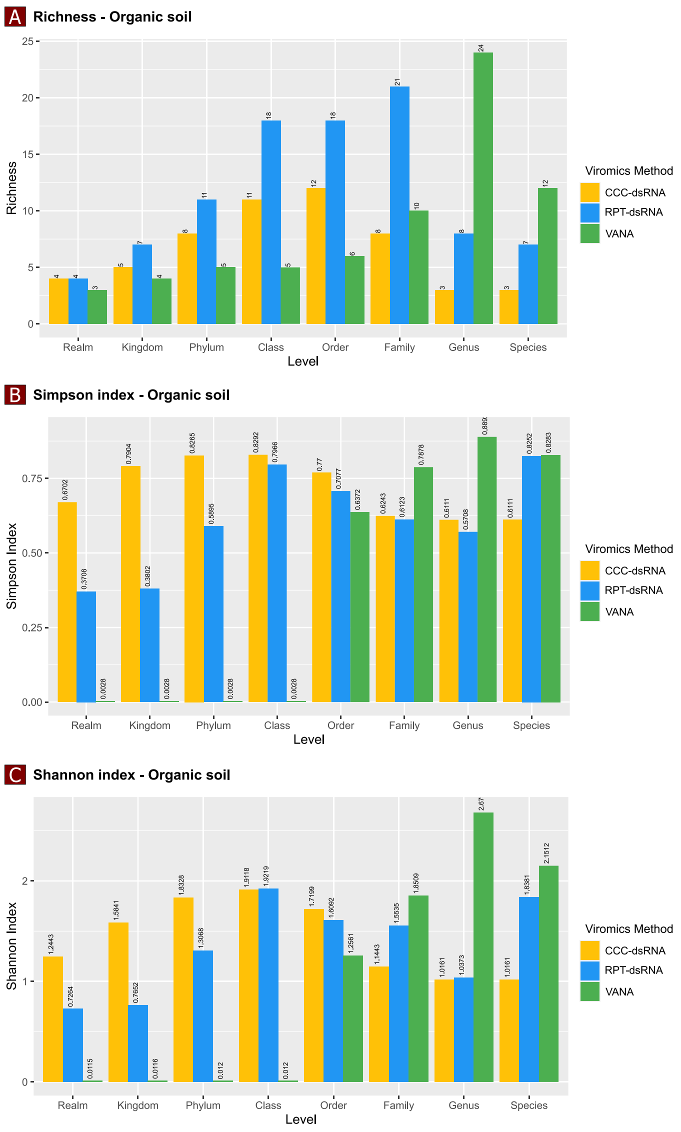


**Supplementary Figure 7: Comparison of alpha-diversity metrics for viral communities in organic soil using three different extraction methods**: Rneasy Power Soil Total RNA kit (RPT-dsRNA), Cellulose-Column Chromatography (CCC-dsRNA), and Virion Associated Nucleic Acid (VANA). The bar chart shows richness (A), Shannon diversity (B), and Simpson diversity (C) indices for each method, calculated using Vegan v2.6-4 and Tidyverse packages. Analysis was performed at the realm, kingdom, phylum, class, order, family, genus, and species levels. The corresponding colors in the legend indicate the different extraction methods used.

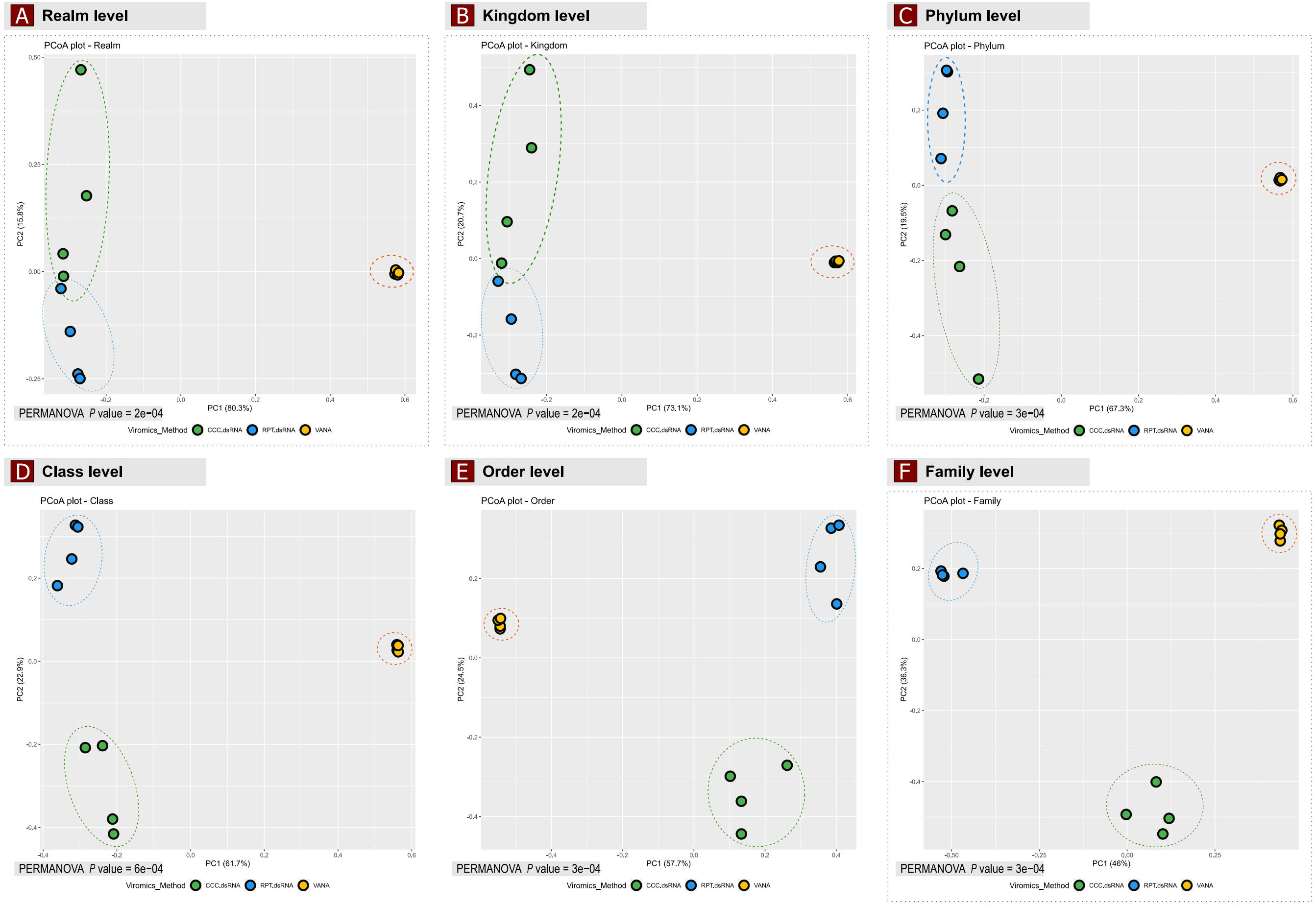


**Supplementary Figure 8: Principal Coordinates Analysis (PcoA) plot of viral community composition in Organic soil, comparing the effectiveness of three extraction methods:** Rneasy Power Soil Total RNA kit (RPT-dsRNA), Cellulose-Column Chromatography (CCC-dsRNA), and Virion Associated Nucleic Acid approach (VANA), based on pairwise Bray-Curtis dissimilarities at each taxonomic rank. Each point on the plot represents a soil sample colored by the extraction method used. The p-value indicates the statistical significance of differences in viral community composition, as determined by a Permutational multivariate analysis of variance (PERMANOVA) test.

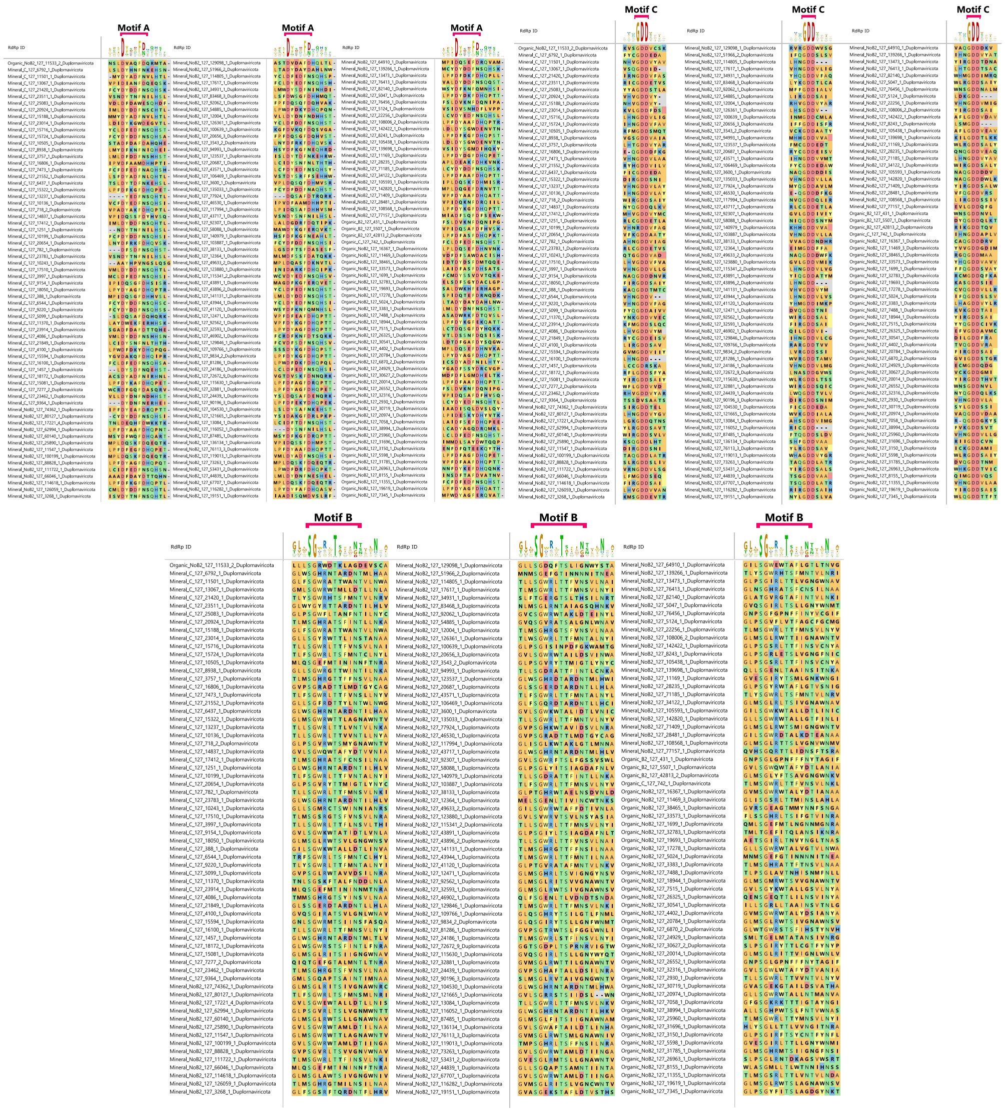


**Supplementary Figure 9:** Multiple sequence alignment (MSA) visualizations of conserved A, B, and C protein motifs within the RdRps domains of novel RdRps in the *Duplornaviricota* phylum (n = 198) using MegAlign Pro v17.4.1 (DNAStar Lasergene software). The MSA was performed to identify conserved regions and potential variations among the candidates of each *Riboviria* taxa. The A, B, and C motif regions are highlighted.

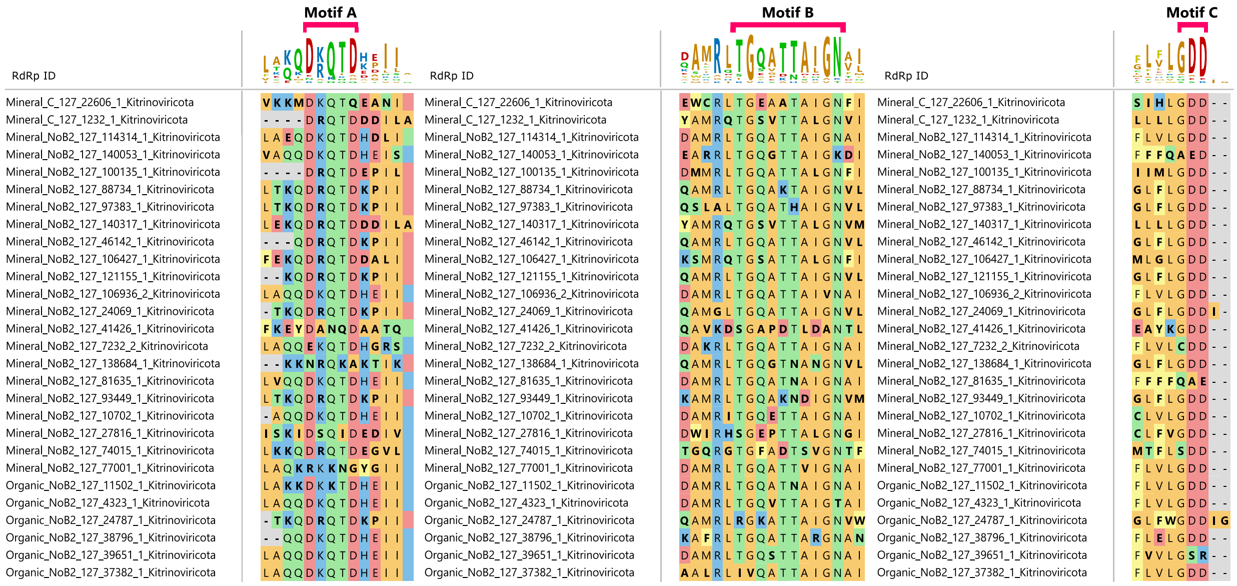


**Supplementary Figure 10:** Multiple sequence alignment (MSA) visualizations of conserved A, B, and C protein motifs within the RdRps domains of novel RdRps in the *Kitrinoviricota* phylum (n =18) using MegAlign Pro v17.4.1 (DNAStar Lasergene software). The MSA was performed to identify conserved regions and potential variations among the candidates of each *Riboviria* taxa. The A, B, and C motif regions are highlighted.

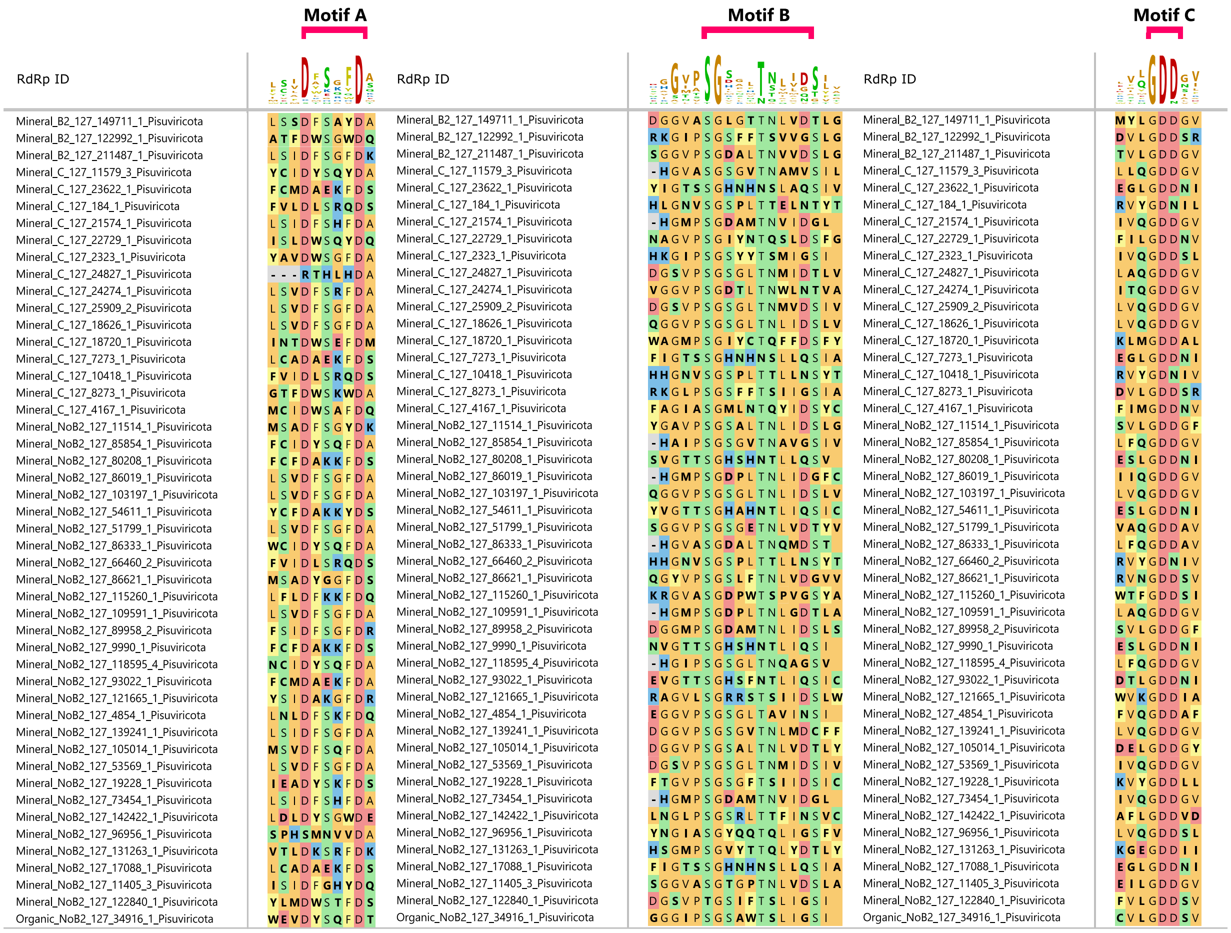


**Supplementary Figure 11:** Multiple sequence alignment (MSA) visualizations of conserved A, B, and C protein motifs within the RdRps domains of novel RdRps in the *Pisuviricota* phylum (n = 48) using MegAlign Pro v17.4.1 (DNAStar Lasergene software). The MSA was performed to identify conserved regions and potential variations among the candidates of each *Riboviria* taxa. The A, B, and C motif regions are highlighted.

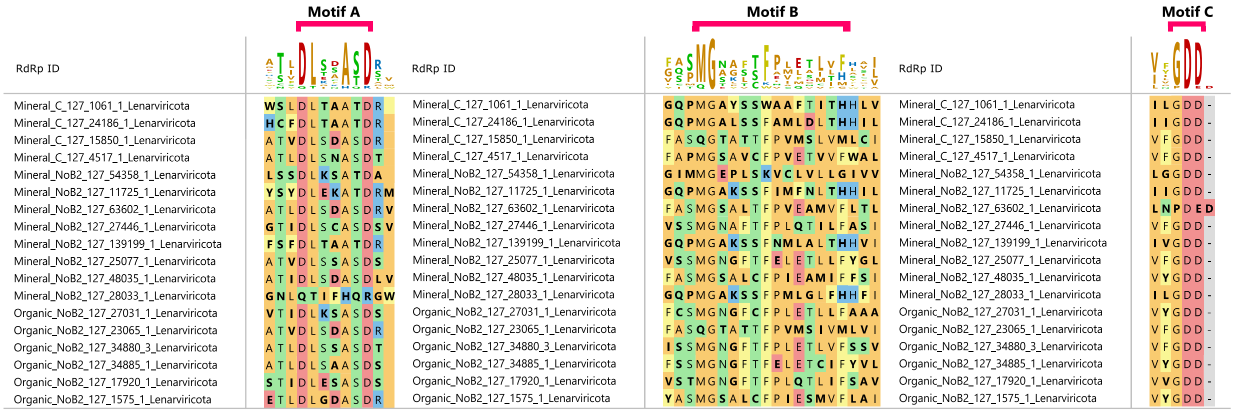


**Supplementary Figure 12:** Multiple sequence alignment (MSA) visualizations of conserved A, B, and C protein motifs within the RdRps domains of novel RdRps in the *Lenarviricota* phylum (n =18) using MegAlign Pro v17.4.1 (DNAStar Lasergene software). The MSA was performed to identify conserved regions and potential variations among the candidates of each *Riboviria* taxa. The A, B, and C motif regions are highlighted.

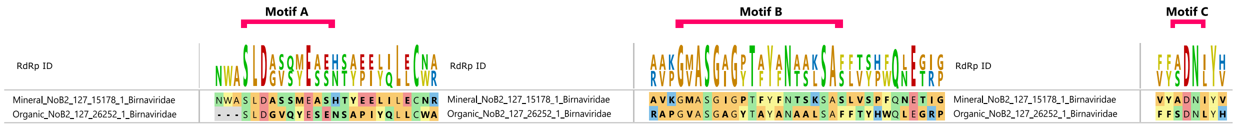


**Supplementary Figure 13:** Multiple sequence alignment (MSA) visualizations of conserved A, B, and C protein motifs within the RdRps domains of novel RdRps in the *Birnaviridae* family (n = 2) using MegAlign Pro v17.4.1 (DNAStar Lasergene software). The MSA was performed to identify conserved regions and potential variations among the candidates of each *Riboviria* taxa. The A, B, and C motif regions are highlighted.
